# Nitazoxanide activates BMP9-ALK1-SMAD signaling cascade and improves HHT vascular pathology

**DOI:** 10.64898/2026.05.12.724733

**Authors:** Camila Chiesa, Valentina Perez-Torrado, Letizia Nada, Rossana Mezzano, Carolina Vazquez, Leonardo Santos, Zelika Criscuolo, Marcelo Serra, Philippe Marambaud, Carlos Escande, Santiago Ruiz

## Abstract

**Objective:** Hereditary hemorrhagic telangiectasia (HHT) is a vascular genetic disorder caused by endothelial cell dysfunction and characterized by telangiectasias and arteriovenous malformations (AVMs). HHT results primarily from loss-of-function mutations affecting components of the BMP9-ALK1-ENG-SMAD signaling cascade, a pathway essential for endothelial quiescence and vascular homeostasis, and currently lacks a cure. Here, we investigated whether nitazoxanide, an orally bioavailable drug with extensive clinical use, can modulate endothelial signaling relevant to HHT.

**Approach and Results:** Nitazoxanide treatment activated SMAD1/5/8 signaling and increased expression of the downstream target ID1 in endothelial cells, while concurrently inhibiting mTOR signaling, indicating a dual modulatory effect on pathways implicated in HHT pathogenesis. In vivo, nitazoxanide activated SMAD signaling in BMP9/10-immunoblocked mice and significantly reduced AVM formation and hypervascularization. Importantly, nitazoxanide restored SMAD1/5/8 activation and ID1 expression in patient-derived blood outgrowth endothelial cells harboring loss-of-function mutations in ALK1 or SMAD4, which exhibit impaired BMP signaling.

**Conclusion:** These findings identify nitazoxanide as a pharmacological modulator capable of activating BMP-SMAD signaling while restraining mTOR activity, thereby overcoming key signaling defects in HHT endothelial cells. Collectively, our results highlight nitazoxanide as a promising therapeutic candidate to target endothelial dysfunction in HHT.

## Introduction

Hereditary hemorrhagic telangiectasia (HHT or Rendu-Osler-Weber syndrome) is an autosomal dominant vascular disease with a prevalence of approximately 1 in 5000 individuals [1]. HHT is characterized by hemorrhagic vascular anomalies in various tissues and organs, manifested as arteriovenous malformations (AVMs). AVMs are direct shunts between arteries and veins lacking an intervening capillary plexus. They become enlarged, tangled, and fragile vessels that depending on their location, can lead to life-threatening complications, including severe epistaxis and internal bleeding, along with secondary complications such as anemia, cerebral abscesses, hepatic issues, pulmonary complications, and cardiac failure [2].

Most HHT patients have loss-of-function mutations in the gene *ENG* (encoding endoglin) or *ACVRL1* (encoding activin receptor-like kinase 1, ALK1), defining the two main subtypes: HHT1 (OMIM #187300) and HHT2 (OMIM #600376), respectively [3,4]. Additionally, mutations in *SMAD4* (encoding SMAD4) and *GDF2* (encoding BMP9) lead to rare forms of the disease called juvenile polyposis/HHT combined syndrome (OMIM #175050) and HHT-like vascular anomaly syndrome (OMIM #615506) [5,6,7]. BMP9, ALK1, endoglin, and SMAD4 functionally interact within the same signaling pathway, comprising the transforming growth factor β (TGF-β) superfamily [8,9]. ALK1, a BMP type I receptor, forms a receptor complex with a BMP type II receptor (e.g., BMPR2) and the co-receptor endoglin. Mutations in BMPR2 can cause familial pulmonary arterial hypertension (PAH), which is observed in some patients with HHT2 [10]. This receptor complex, abundantly expressed in endothelial cells (ECs), is specifically activated by the circulating ligands BMP9 and BMP10 [11–14]. Upon activation, ALK1 receptor phosphorylates the signal transducers SMAD1, SMAD5, and SMAD8, leading to the formation of SMAD1/5/8–SMAD4 complexes that translocate to the nucleus to regulate specific gene expression programs [15–17].

Loss-of-function mutations in ENG and ACVRL1 hinder Smad1/5/8 signaling by BMP9 [18–20], which contributes to the vascular anomalies associated with HHT, such as vascular hyperproliferation and AVMs [21–24]. While the precise cellular processes underlying AVM development in HHT - characterized by direct shunts between arteries and veins - remain poorly understood, evidence indicates that HHT arises from abnormal reactivation of angiogenesis [25,26]. Through angiogenesis, ECs engage distinct transcriptional programs that drive changes in morphology, proliferation, migration, and sprout assembly to form new vascular structures [27]. This process is initiated by vascular endothelial growth factor (VEGF) signaling through VEGF receptor 2 (VEGFR2) in a subset of ECs, leading to the disruption of endothelial quiescence and the reprogramming of cellular homeostasis. These changes alter intercellular signaling dynamics and promote the initiation of new vessel sprouts. Within this context, ALK1 signaling integrates with VEGF-driven pathways to fine-tune endothelial responses, reinforce transcriptional programs that maintain cellular specialization, and ensure balanced vessel growth. Through these regulatory functions, ALK1 acts as a critical regulator of vascular development and homeostasis [24,27].

Mechanistic studies show that ACVRL1 silencing enhances VEGFR2 phosphorylation and signaling [28], while ENG silencing alters VEGFR2 trafficking [29], both contributing to excessive pathway activation. Transcriptomic analyses further demonstrate that ALK1 negatively regulates VEGFR2 expression [30, 31], and *in vivo* studies reveal that genetic or pharmacological inhibition of VEGFR2 reduces hypervascularization and AVMs in HHT models [28, 29]. Additionally, and more recently, overactivation of the PI3K/AKT/mTOR pathway has been identified as a key driver of vascular pathology in HHT. In ALK1- and endoglin-deficient models, ECs show increased phosphorylation of ribosomal protein S6, a hallmark of mTOR activation, and blocking PI3K or AKT reduces AVM formation [28,29,31]. We found that sirolimus potently inhibits mTOR signaling while also activating SMAD1/5/8 via ALK2, thereby correcting two major defects in HHT ECs. In mouse models, sirolimus treatment normalized mTOR activity, reduced AVM burden, and improved vascular pathology at clinically relevant doses [32]. These findings identify mTOR as a central pathogenic node in HHT and support mTOR inhibition as a promising therapeutic approach.

In recent years, multiple pharmacological strategies have been explored for HHT treatment. These include, among others, inhibitors of VEGF (bevacizumab), PI3K (pictilisib), the receptor tyrosine kinase targeting VEGFR2 (pazopanib), and CDK4/6 (palbociclib and ribociclib), and activators of SMAD1/5/8 (tacrolimus and sirolimus), alone or in combination. Several of these agents have shown beneficial effects in preclinical models, particularly reducing the number of AVMs [33–36]. More importantly, clinical trials of some of them have demonstrated therapeutic benefit, especially in reducing HHT-associated epistaxis [37–42]. As trials develop, knowledge allows us to make considerations and tweak approaches to improve the drug’s efficacy, effects, adequacy to its personalized usage considering health condition, treatment incompatibilities and known adverse events, and lastly to increase the drug tool kits for HHT therapeutics.

In this study, we investigated whether the FDA-approved drug nitazoxanide, a drug that we identified as a possible activator of endothelial SMAD1/5/8 signaling [31], could potently activate the signaling pathway to prevent vascular anomalies and pathology in HHT. We found that nitazoxanide is a potent SMAD1/5/8 signaling activator, producing effects comparable to tacrolimus and sirolimus in ECs. Additionally, nitazoxanide restored SMAD1/5/8 signaling through ALK1 activation and prevented vascular anomalies and HHT pathology in the BMP9/BMP10-immunoblocked (BMP9/10ib) mouse model. Furthermore, nitazoxanide activated SMAD1/5/8 signaling in blood outgrowth endothelial cells (BOECs) derived from patients with HHT2 and HHT/JPS, supporting its therapeutic potential in HHT.

## Materials and Methods

### Reagents and antibodies

Recombinant human BMP9 (#3209-BP) and VEGF (#293-VE) were obtained from R&D Systems. Nitazoxanide (#24609) was from AstaTech, Tacrolimus (#10007965) from Cayman Chemicals and Sirolimus (#J62473) from Thermo Fisher Scientific. For Western blots, antibody directed against pSMAD1/5/8 (#13820), anti-SMAD1 (#6944), anti-AKT (#9272), anti-pS473-AKT (#4060), anti-pS6 (#2211), anti-S6 (#2217), anti-ID3 (#9837), anti-pp38 (#4511) and anti-p38 (#8690) were obtained from Cell Signaling Technology; anti-ID1 antibody (#BCH-1/195-14) from BioCheck; anti-β-actin antibody (#A5441) and anti-ALK2 antibody (#155981) from Abcam; anti-ALK1 antibody (#PA5-14921) from Invitrogen. Goat anti-rabbit IgG secondary antibody, HRP-conjugate (#A0545) and rabbit anti-mouse IgG secondary antibody, HRP-conjugate (#A9044) were obtained from Sigma Aldrich. For protein concentration, Pierce BCA Protein Assay Kit (#23225) was obtained from Thermo Fisher Scientific. For protein extraction, cOmplete™, EDTA-free protease inhibitor cocktail tablets (COEDTAF-RO) was obtained from Roche and zirconium oxide beads (ZrOB05) from Next Advance. Bovine serum albumin (BSA; BSA-1U) was obtained from Capricorn Scientific. For immunoblocking of BMP9 and BMP10 in mice, anti-BMP9 (#MAB3209) and anti-BMP10 (#MAB2926) were obtained from R&D Systems. For flow cytometry, CD45-APCCy7 antibody (#103115) and Zombie Aqua fixable viability kit (#423101) were obtained from BioLegend, pSMAD1/8-PE antibody (#562509) was obtained from BD Biosciences, and eBioscience^TM^ FOXP3/Transcription Factor staining buffer set (#00552300) was from Invitrogen. Alexa Fluor 546-conjugated goat anti-rabbit (#A11035), Alexa Fluor 488 goat anti-rabbit (#A11034), Alexa Fluor 488 goat-anti rat (#A11006), TO-PRO3 (#S33025) and isolectin GS-IB_4_ conjugated to Alexa Fluor 488 (#21411) were obtained from Invitrogen, DAPI (#D9542) from Sigma Aldrich, and anti-CD31 (#550274) from BD Biosciences. ProLong Gold antifade reagent (#P36930) was obtained from Invitrogen. For tissue embedding and cryopreservation, Tissue-Tek® O.C.T. Compound (4583) was obtained from Sakura Finetek. Stealth siRNAs targeting ACVRL1 and ACVR1, and a negative control, were obtained from Invitrogen. Lipofectamine RNAiMAX Transfection Reagent (#13778-075) was obtained from Invitrogen. LDN193189 (#6053) was obtained from Tocris Bioscience. qPCR primers targeting *ACVRL1* and *ACVR1,* and Oligo-dT primers were purchased from Integrated DNA Technologies (IDT). FastStart Universal SYBR Green Master (#04913914001) was obtained from Roche and SuperScript II Reverse Transcriptase (#18064014) was obtained from Thermo Fisher Scientific.

### Cells

HUVECs were obtained from Lonza and were cultured in EC growth medium (R&D Systems) supplemented with penicillin and streptomycin. C2C12 cells were obtained from ATCC and were maintained in Dulbecco’s Modified Eagle’s Medium (DMEM) supplemented with 10% FBS, penicillin and streptomycin. Blood outgrowth endothelial cells (BOECs) were isolated from blood draws obtained from HHT patients. Cells were isolated as described before [43].

### Cell treatments

HUVECs treated or not for 24 hours in complete medium (conditioned for 2 days), with 1 µM Tacrolimus, 1 µM Sirolimus, 1 µM Nitazoxanide or 10 ng/mL BMP9 were stimulated or not for 5 min with 25 ng/mL VEGF. C2C12 and HUVECs were treated or not for 24 hours in complete medium (conditioned for 2 days), with different concentrations of the drug (0.03, 0.1, 0.3, 1 µM). BOECs from six HHT patients were treated or not for 24 hours with 1 µM nitazoxanide.

### Flow cytometry

Flow cytometry was performed to evaluate the effect of 0.3 and 1 µM nitazoxanide in C2C12 cells and to characterize the generated BOECs, using CD45 as a hematopoietic (negative) marker. C2C12 cells were treated with nitazoxanide as previously described and stained with Zombie Aqua (1:500) for viability and anti-pSMAD1/8-PE antibody (1:50). Samples were acquired using a Cytek Aurora spectral flow cytometer. Cells were first identified based on morphology (SSC-A vs FSC-A), followed by singlet discrimination (FSC-A vs FSC-H), and gating of live cells (Zombie–). The percentage of live cells was determined, and within this population, the percentage of pSMAD1/8-positive cells was analyzed. Positivity for pSMAD1/8 was defined using fluorescence minus one (FMO) controls for each condition. BOECs were stained with Zombie Aqua (1:500), and anti-CD45 antibody (1:200). Data acquisition was performed using an Attune NxT cytometer. Cells were first identified based on morphology (SSC-A vs FSC-A), followed by singlets selection (FSC-A vs FSC-H), and gating of live cells (Zombie–). Within the live cell population, BOECs were defined as CD45-negative cells, confirming the successful isolation and expansion of this population during cell culture. Data analysis was performed using FlowJo v10.8.

### Small interfering RNA

Pharmacological inhibition was performed using the pan-ALK inhibitor LDN193189 in HUVECs. Cells were cultured for two days in complete medium and treated or not for 24 hours with 120 nM LDN193189 and 1 µM Nitazoxanide. RNA interference (RNAi) was performed in HUVECs using Stealth RNAi siRNAs targeting *ACVRL1* (20 nM) and *ACVR1* (20 nM), or using a stealth negative control (20 nM), and by following the manufacturer’s recommended protocols (Invitrogen). The effectiveness of the specific siRNAs was confirmed 48 hours later by qPCR and Western blot. The silenced cells were treated for 24 hours with 1 µM Nitazoxanide, and pSMAD1/5/8 and ID1 levels were determined by Western blot.

### RT-qPCR

Two days after siRNA treatment, cells were harvested and total RNA was extracted using the TRIzol–chloroform method. RNA concentration and purity were assessed using a NanoDrop spectrophotometer (Thermo Fisher Scientific). Complementary DNA (cDNA) was synthesized from 2 µg of total RNA using 100 U of oligo(dT) primers and SuperScript II Reverse Transcriptase, according to the manufacturer’s instructions (Invitrogen). Quantitative PCR (qPCR) was performed using primers targeting ACVRL1, ACVR1 and B2M (housekeeping) [44]. Amplification was carried out in a QuantStudio 3 system (Applied Biosystems). Relative changes in gene expression were determined by the ΔΔCt method.

### Immunoblocking of BMP9 and BMP10, nitazoxanide treatments and retinal vasculature analyses in mice

C57BL/6 mouse pups were administered anti-BMP9 (12.5 mg/kg) and anti-BMP10 (25 mg/kg) antibodies via intraperitoneal (i.p.) injection on postnatal days 3 (P3) and 4 (P4). On P3, P4 and P5, pups were injected i.p. with nitazoxanide (10 and 20 mg/kg) or vehicle (DMSO). Pups were euthanized on P6 by CO_2_ asphyxiation and eyes, liver and lungs were collected. Eyes were fixed in 4% PFA for 30 minutes on ice and retinas were dissected, cut four times to allow them to be laid flat and fixed with methanol for 20 minutes on ice. After removing methanol, retinas were rinsed in PBS for 10 minutes on a shaker at RT and incubated in blocking solution (0.3% Triton, 0.2% BSA, 10% normal goat serum in PBS) for 1 hour at RT. For staining, retinas were incubated overnight at 4°C with isolectin GS-IB_4_ conjugated to Alexa Fluor 488 (1:100, 0.3% Triton in PBS) and pSMAD1/5/8 (1:1000, 0.3% Triton in PBS). Retinas were washed four times with 0.3% Triton in PBS. When only isolectin GS-IB_4_ was used, retinas were washed again with PBS for 10 minutes and then mounted onto slides using ProLong Gold antifade reagent. For pSMAD1/5/8 immunofluorescence, retinas were incubated with goat anti-rabbit IgG conjugated to Alexa Fluor 546 (1:1000, 0.3% Triton in PBS) for 2 hours at RT. Retinas were then washed three times with 0.3% Triton in PBS and once with PBS for 10 minutes before mounting. Retinas were imaged on an Olympus IX81 epifluorescence microscope at 4x, 10x and 20x magnifications and on a Zeiss LSM800 confocal microscope at 63x magnification. Quantifications were performed using ImageJ v1.54k software. Images were acquired in three different locations of the vasculature: the plexus (between a vein and an artery) using a 10x lens, and the front of the extending arterial and venous vasculature using a 20x lens. The area occupied by the vasculature in a region of interest of 100×100 µm^2^ was measured in these three locations. For the analysis of the number and diameter of AVMs, and the diameter, whole retinas were imaged with a 4x lens.

### Western blot

After the experiments described above, cells were washed with PBS. C2C12 and HUVECs were collected in a 1x loading buffer. BOECs were collected with RIPA buffer containing 5 mM sodium fluoride, 50 mM β-glycerophosphate and 1x cOmplete, EDTA-free protease inhibitor cocktail (Roche), quantified using BCA method and prepared for western blot analysis with 3x loading buffer. Tissues (lung and liver) were homogenized in RIPA buffer supplemented with 5mM sodium fluoride, 50 mM β-glycerophosphate and 1x cOmplete, EDTA-free protease inhibitor cocktail (500 µL of RIPA per 50 mg of frozen tissue). A small spoon full of zirconium oxide beads was added, and homogenization was performed using a Bullet Blender for 3 minutes at speed 10. Lysates were centrifuged at 12,000 x g for 10 minutes at 4°C. The supernatants were collected, and protein concentration was determined using the BCA assay. Samples were separated by SDS-PAGE and transferred onto nitrocellulose membranes. Membranes were then probed with primary and secondary antibodies. A standard ECL detection procedure was then used. Protein expression was quantified by densitometry using ImageJ v1.54k software, and relative expression was calculated as the ratio of the signal of the protein of interest to actin or the corresponding total protein.

### Immunocytochemistry and immunohistochemistry

C2C12 cells were plated onto 0.5% gelatin-coated coverslips (10 mm) and cultured in complete EC growth medium for two days. Cells were treated or not for 24 hours with different concentrations of the drug (0.03, 0.1, 0.3 and 1 µM), washed with PBS, fixed in 4% PFA for 20 minutes, and then permeabilized in 0.3% Triton x-100 in PBS (PBS-T). After blocking with 10% normal goat serum in PBS-T for 1 hour at room temperature, cells were incubated overnight at 4°C with rabbit anti-pSMAD1/5/8 antibody (1:1000), diluted in PBS-T. Cells were then washed and incubated for 2 hours at room temperature with Alexa Fluor 546-conjugated goat anti-rabbit secondary antibody, along with DAPI, all diluted in PBS-T. Cells were mounted on ProLong Gold antifade reagent and imaged under the microscope. For tissue imaging, lungs and liver were fixed in 4% PFA overnight at 4°C, washed once with PBS, and transferred to PBS containing 15% sucrose for overnight incubation at 4°C. The solution was then replaced with PBS containing 30% sucrose before immersing the tissues in OCT compound and freezing for cryosectioning. Sections (10-12 µm) were mounted on slides and permeabilized with 0.3% Triton in PBS for two consecutive 10-minute incubations at RT. After permeabilization, the sections were blocked for 1 hour at RT in 0.3% Triton, 0.2% BSA, 5% normal goat serum in PBS. Antibodies against pSMAD1/5/8 (1:1000) and pS6 (1:1000) were then applied in 0.3% Triton in PBS for lungs and liver sections, respectively, and the slides were incubated for 48 hours at 4°C in a humidified chamber without agitation. The sections were washed three times for 10 minutes in 0.3% Triton in PBS, followed by a 2-hour incubation at RT with goat anti-rabbit IgG conjugated to Alexa Fluor 546 (1:1000) and DAPI (1:1000) prepared in 0.3% Triton in PBS. Stained sections were washed three times for 10 minutes in 0.3% Triton in PBS and twice for 5 minutes in PBS, and then mounted using ProLong Gold antifade reagent. In both cases, imaging was performed using a Zeiss LSM800 confocal microscope at 63x magnification.

### Statistical Analysis

For all datasets, normality was assessed using the D’Agostino–Pearson omnibus test or the Shapiro–Wilk test, as appropriate. Homogeneity of variances (homoscedasticity) was evaluated using the Brown–Forsythe test for multiple-group comparisons and the F test for two-group comparisons. For multiple-group comparisons, when data fulfilled both normality and homoscedasticity assumptions, a one-way ANOVA followed by Tukey’s multiple-comparisons test was performed. When data were normally distributed but did not meet the assumption of equal variances, Welch’s ANOVA followed by Dunnett’s T3 multiple-comparisons test was used. When normality was not achieved, a nonparametric Kruskal–Wallis test followed by Dunn’s multiple-comparisons test was applied. For two-group comparisons, when data fulfilled both normality and homoscedasticity assumptions, a Student’s t-test was used. When data were normally distributed, but variances were unequal, a t-test with Welch’s correction was applied. When data were not normally distributed, a Mann–Whitney test was performed. For two-way experimental designs, data were analyzed using two-way ANOVA followed by Tukey’s multiple-comparisons test when parametric assumptions were met. Homoscedasticity for these analyses was assessed using Levene’s test. When normality or homoscedasticity were not achieved, data were log-transformed to satisfy these assumptions before analysis. In the case of experiments with BOECs, paired statistical analyses were applied. When data fulfilled normality assumptions, a paired Student’s t-test was used. When normality was not achieved, a Wilcoxon matched-pairs signed-rank test was performed instead. *P* values less than 0.05 were considered statistically significant. Analyses were done using GraphPad Prism, except for normality and homoscedasticity tests associated with two-way ANOVA, which were conducted in RStudio.

### Study approval

Study subjects (HHT2 and HHT/JPS patients carrying a mutation in SMAD4) provided voluntary and written informed consent using a form approved by Centro Hospitalario Pereira Rossell Institutional and Hospital Italiano de Buenos Aires Review Board (IRB). Study subject BOECs were isolated and cultured using a protocol approved by both Institutions’ IRB. Clinical trial number: not applicable. All animal procedures were performed in accordance with protocols (009-24) approved by the Institut Pasteur de Montevideo Institutional Animal Care and Use Committees.

## Results

### Nitazoxanide is a potent SMAD1/5/8 signaling activator

In previous work, we identified immunosuppressants tacrolimus and sirolimus as potent activators of SMAD1/5/8 signaling [31,32], a key downstream effector of the BMP9-ALK1-SMAD signaling cascade. Consistent with this, clinical reports have shown that both drugs can reduce severe epistaxis and gastrointestinal bleeding in patients with HHT [45–49]. However, given their immunosuppressive nature and the need for lifelong treatment in HHT, we took a step forward in developing a safer long-term therapeutic drug that restores the BMP9-ALK1-SMAD cascade while avoiding chronic immunosuppression and expanding patient eligibility. To this end, we focused on nitazoxanide, another promising candidate previously identified through screening the NIH clinical collections for activators of BMP9-specific SMAD1/5/8 signaling [31]. We first compared the effect of nitazoxanide to tacrolimus and sirolimus on SMAD1/5/8 signaling. HUVECs were cultured in complete medium containing the serum’s exogenous trophic factors, conditioned for two days, and then treated with each drug at 1 µM for 24 hours. Western blot analysis showed that nitazoxanide efficiently increased SMAD1/5/8 phosphorylation and induced ID1 (inhibitor of differentiation 1) expression at the protein level, with an efficacy comparable to tacrolimus and sirolimus (Fig. 1). Relative to untreated controls, both pSMAD1/5/8 and ID1 levels were significantly elevated. These results suggest that nitazoxanide is a potent trigger of the BMP9-ALK1-SMAD signaling cascade in endothelial cells.

**Figure 1.**
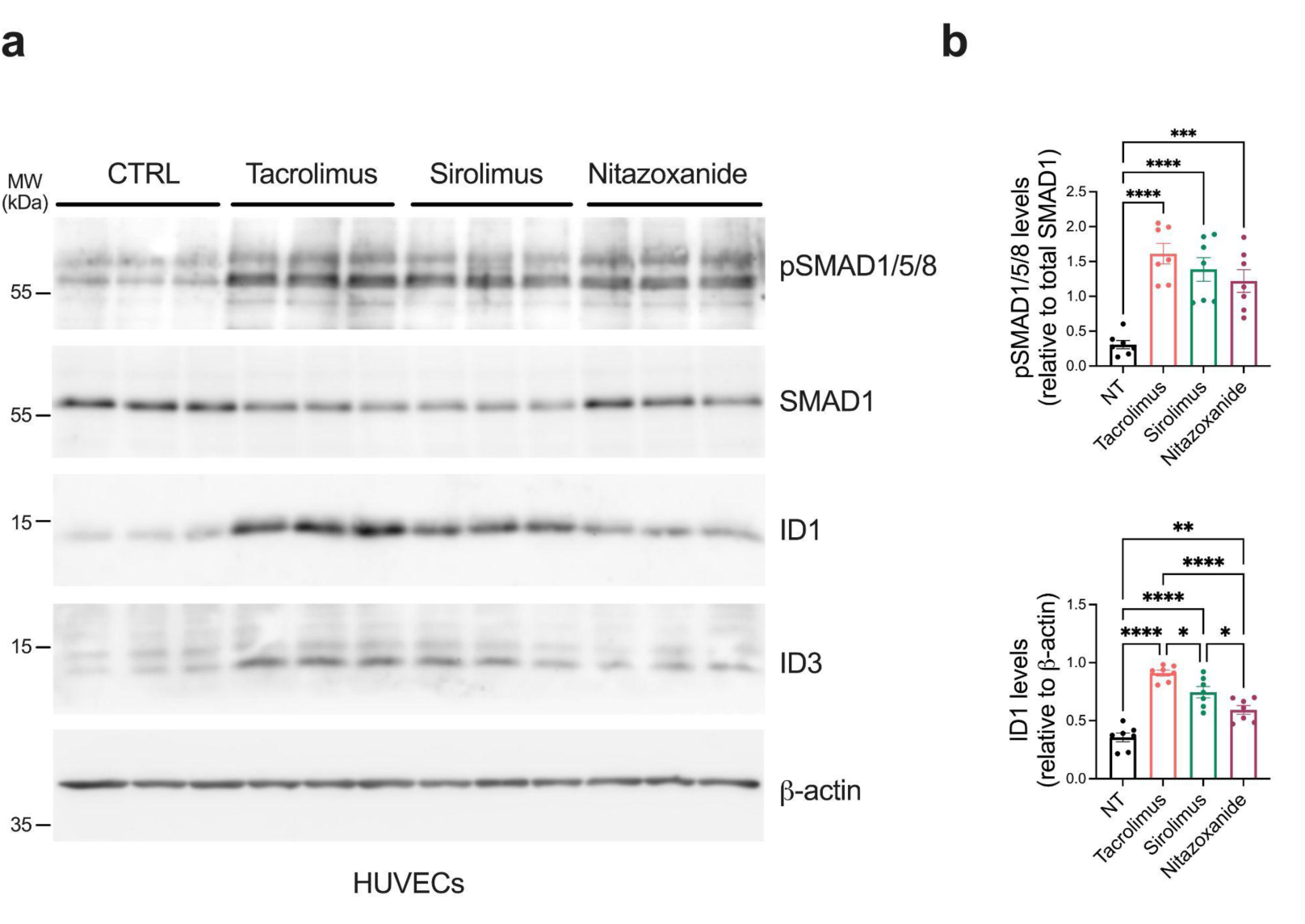
Nitazoxanide activates the BMP9-ALK1-SMAD signaling pathway with efficacy comparable to that of the immunosuppressants tacrolimus and sirolimus. HUVECs were treated or not (NT) for 24 hours in complete medium (conditioned for 2 days) with 1 μM tacrolimus, 1μM sirolimus or 1 μM nitazoxanide. (a) Cell extracts were analyzed by Western blot using antibodies directed against the indicated proteins. (b) Densitometric analyses and quantification of pSMAD1/5/8, ID1 and ID3 relative levels in n=3 independent experiments. Data represent mean ± SEM; **p<0.01, ***p<0.001, ****p<0.0001, one-way ANOVA followed by Tukey’s multiple-comparisons test.

To further characterize the effect of nitazoxanide on SMAD1/5/8 signaling, dose-response experiments were performed in both C2C12 cells (Fig. 2a) and HUVECs (Fig. 2b). Cells were cultured in complete medium, conditioned for two days, and then treated with increasing concentrations of nitazoxanide (0.03, 0.1, 0.3 and 1 µM) for 24 hours. Western blot analysis revealed that nitazoxanide increased SMAD1/5/8 phosphorylation in a dose-dependent manner and also increased ID1 protein expression levels (Fig. 2a-d). In addition, confocal microscopy of cells stained for pSMAD1/5/8 and DAPI for nuclear staining clearly showed an increase in nuclear pSMAD1/5/8 fluorescence (Fig. 2e-f). Flow cytometry analysis of C2C12 cells treated with 0.3 and 1 µM of nitazoxanide further supported these findings, consistently confirming that nitazoxanide increases pSMAD1/5/8 levels. Both concentrations increased pSMAD1/8 levels, with a higher percentage of phosphorylation at 1 µM (Suppl. Fig. 1a-b), consistent with Western blot results (Fig. 2a-d). Additionally, no changes in cell viability were detected (Suppl. Fig. 1c-d). Together, these data identify nitazoxanide as a potent activator of SMAD1/5/8 signaling.

**Figure 2.**
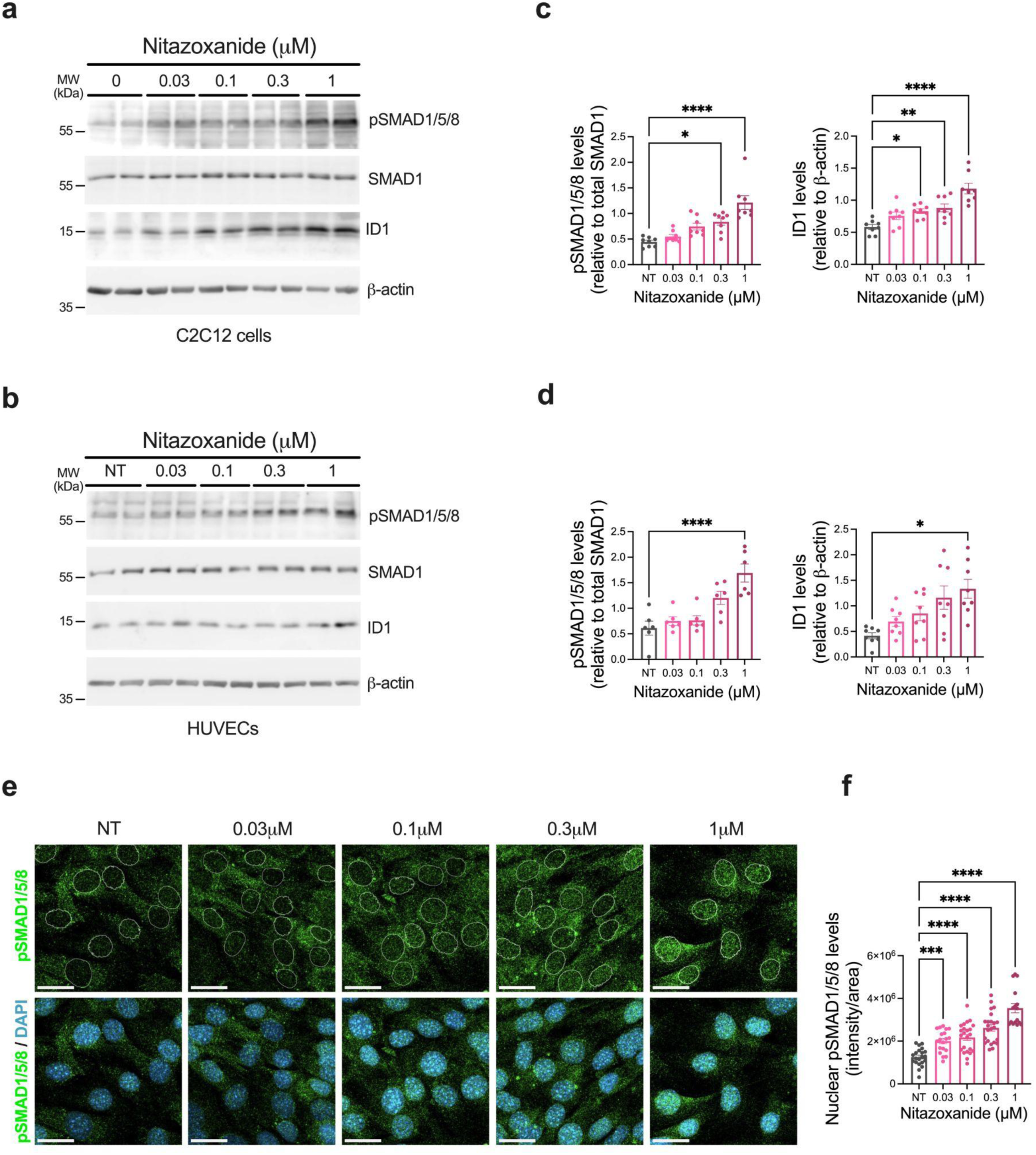
Nitazoxanide is a potent SMAD1/5/8 signaling activator. (a) C2C12 cells and (b) HUVECs were treated or not (NT) for 24 hours in complete medium (conditioned for 2 days) with nitazoxanide at the indicated concentrations. Cell extracts were analyzed by Western blot using antibodies directed against the indicated proteins. (c-d) Densitometric analyses and quantification of pSMAD1/5/8 and ID1 relative levels in n=3 independent experiments. Data represent mean ± SEM; *p<0.05, **p<0.01, ****p<0.0001, one-way ANOVA followed by Tukey’s multiple-comparisons test (c, ID1; d, pSMAD1/5/8); Kruskal-Wallis followed by Dunn’s multiple-comparisons test (b, pSMAD1/5/8) and one-way ANOVA followed by Dunnett’s T3 multiple-comparisons test (d, ID1). (e) Representative immunofluorescence (IF) confocal images of C2C12 cells treated for 24 hours with nitazoxanide at the indicated concentrations. Cells were labeled with DAPI (blue) and pSMAD1/5/8 (green). Scale bars, 25 μm. (f) Nuclear pSMAD1/5/8 fluorescence intensity quantification. ***p<0.001, ****p<0.0001, one-way ANOVA followed by Dunnett’s T3 multiple-comparisons.

### Nitazoxanide rescues SMAD1/5/8 signaling by activating ALK1

To elucidate the mechanism by which nitazoxanide activates the BMP9-ALK1-SMAD signaling cascade, we combined pharmacological inhibition with genetic silencing approaches in HUVECs (Fig. 3 and Suppl. Fig. 2). Cells were cultured for two days and then treated for 24 hours with the pan-ALK inhibitor LDN193189 (120 nM) and/or 1 µM nitazoxanide. Under ALK inhibition, nitazoxanide failed to increase ID1 expression or SMAD1/5/8 phosphorylation (Fig. 3a-b), indicating that its effect depends on ALKs’ activation.

**Figure 3.**
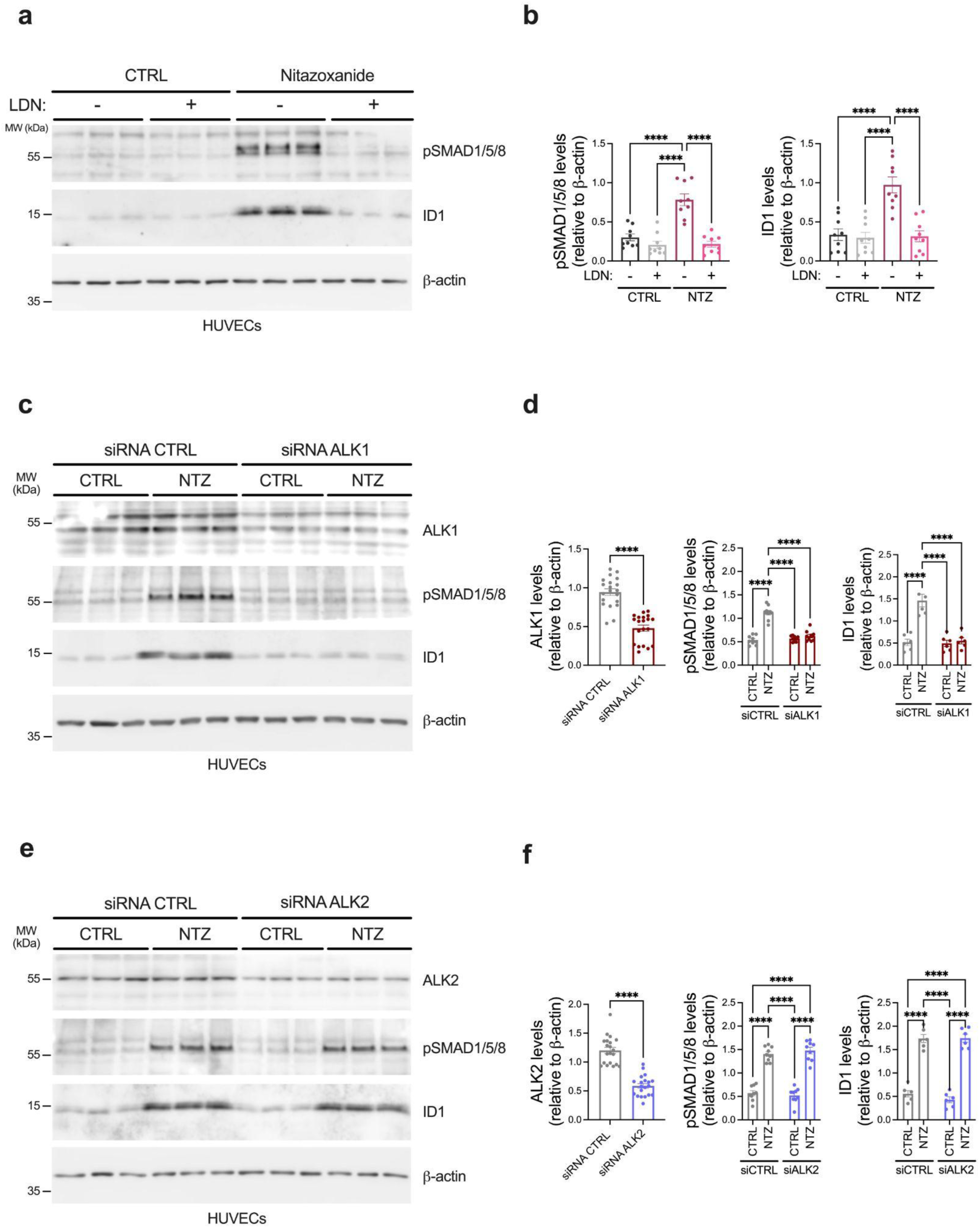
Nitazoxanide activates SMAD1/5/8 via ALK1. Effect of ALK1 and ALK2 silencing on nitazoxanide-induced activation of the BMP9-ALK1-SMAD signaling pathway in HUVECs. (a) HUVECs were treated for 24 hours with 1 μM nitazoxanide in the absence or presence of LDN193189 (LDN, 120 nM). Cell extracts were analyzed by Western blot using antibodies directed against the indicated proteins. (b) Densitometric analyses and quantification of pSMAD1/5/8 and ID1 relative levels in n=3 independent experiments. Data represent mean ± SEM. ****p<0.0001, one-way ANOVA followed by Tukey’s multiple-comparisons test. (c,e) HUVECs were transfected with 20 nM Stealth siRNAs targeting (c) *ACVRL1* (ALK1) and (e) *ACVR1* (ALK2), or a Stealth RNAi negative control (20 nM), followed by treatment with 1 μM nitazoxanide overnight. Phosphorylation of SMAD1/5/8 and ID1 expression levels were assessed by Western blot. (d,f) Densitometric analyses and quantification of (d) ALK1, (f) ALK2, and (d,f) pSMAD1/5/8 and ID1 relative levels in n=3 independent experiments. Data represent mean ± SEM. ****p<0.0001, unpaired two-tailed Student’s t-test (d, ALK1; f, ALK2) and two-way ANOVA followed by Tukey’s multiple-comparisons test (pSMAD1/5/8 and ID1).

To distinguish between the involvement of specific ALK receptors, our silencing approach targeted *ACVRL1* (which encodes ALK1), *ACVR1* (which encodes ALK2) and a negative control to assess non-specific effects in HUVECs. Knockdown efficiency for both genes was analyzed at protein (Fig. 3c-f) and mRNA levels (Suppl. Fig. 2), confirming that siRNAs significantly reduced their intended targets’ expression. These silenced cells were then treated for 24 hours with 1 μM nitazoxanide to determine whether the drug rescues BMP9 signaling differentially in ALK1- or ALK2-silenced cells as observed in control cells. Subsequently, western blot analysis revealed that pSMAD1/5/8 and ID1 expression were not induced in ALK1-silenced cells (Fig. 3c-d), whereas a clear increase in both markers was observed in ALK2-silenced cells (Fig. 3e-f). These results demonstrate that nitazoxanide activates the BMP9-ALK1-SMAD signaling cascade through the ALK1 receptor.

### Nitazoxanide prevents vein dilation, hypervascularization and AVM formation in the retina of BMP9/BMP10-immunoblocked mice

To gain insight into the therapeutic potential of nitazoxanide *in vivo*, we used the neonatal BMP9/10ib model. This model reproduces key features of HHT in the retinal vasculature, including venous dilation, excessive vascular branching, and AVM formation [31, 32]. Before efficacy studies, dose-finding experiments were performed to identify the highest nitazoxanide regimen and dose that did not alter normal retinal vascular development (Suppl. Fig. 3a-d) but that generated a slight reduction in the vascular density in wild type (CTRL) mice. Notably, we found that the highest tested concentrations (20 mg/kg/day) significantly reduced the vascular density of all analyzed areas, vascular plexus (Suppl. Fig. 3e-h), and front of veins (Suppl. Fig. 3i-l) and arteries (Suppl. Fig. 3m-p). We also used this dose-finding experiment (Fig. 4a) to evaluate pS6 and pSMAD1/5/8 expression levels in liver and lungs, respectively. With this approach, we defined that nitazoxanide significantly decreased pS6 levels in the liver (Fig. 4b-c), and increased pSMAD1/5/8 levels and ID1 expression in the lungs of wild type (CTRL) mice (Fig. 4d-e). Based on these results, BMP9/10-immunoblocked pups were treated preventively by daily i.p. injection with nitazoxanide (20 mg/kg/day) from P3 to P5 (Fig. 4f). Mice were then analyzed at P6, a time point at which vascular abnormalities, including vessel dilation, hypervascularization, and AVMs, can be identified and quantified. Nitazoxanide treatment markedly attenuated these abnormalities induced by BMP9/10 immunoblocking (Fig. 4g-w). Nitazoxanide significantly reduced both the number and size of AVMs (Fig. 4j-k, l-n). In addition, nitazoxanide significantly decreased venous dilation (Fig. 4o). Quantitative analyses revealed that nitazoxanide decreased the BMP9/10 immunoblocked-dependent increase in density of the vascular plexus (Fig.4 g-i, p-r and v) and the front of artery (Fig.4 g-i, s-u and w), reflecting the prevention of vascular abnormalities in the BMP9/10ib retinas. Together, these data demonstrate that nitazoxanide protects against HHT-like retinal vascular pathology *in vivo*. The drug effectively restrained hypervascularization and venous dilation, and reduced AVM formation, supporting its potential as a candidate for further preclinical evaluation in the context of HHT.

**Figure 4.**
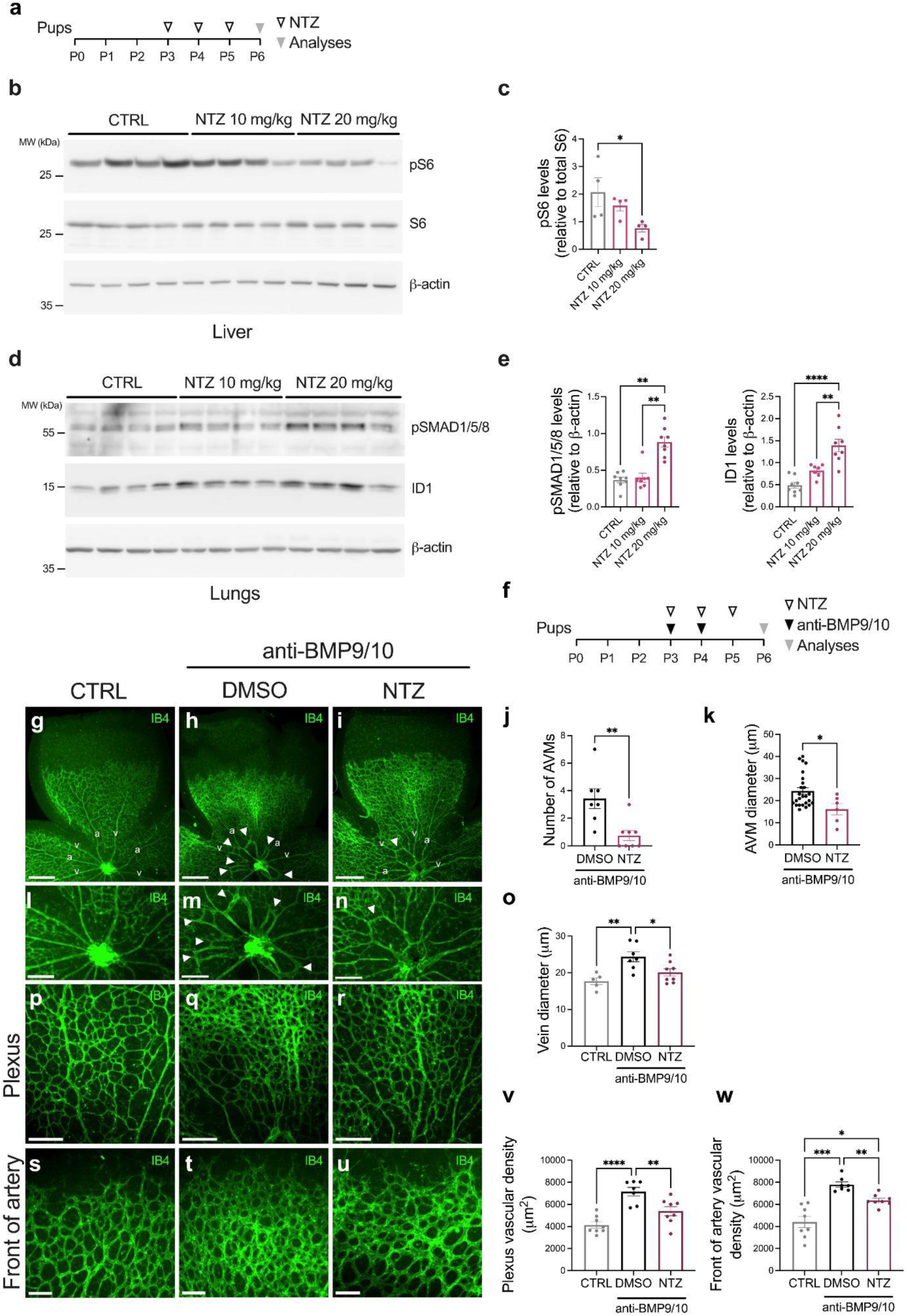
Nitazoxanide exerts dose-dependent effects on the liver and lung of wild type mice and prevents vein dilation, hypervascularization, and AVMs in the BMP9/10ib mouse retina. (a) Experimental strategy of nitazoxanide dose-finding approach in wild type (CTRL) mice. Pups were injected i.p. from P3 to P5 with 10 mg/kg/d or 20 mg/kg/d nitazoxanide. Arrowheads indicate the timepoints of injections and analyses. Pups were euthanized on P6 for analysis. (b, d) Western blot analysis of (b) liver or (d) lung tissue homogenates from P6 pups injected at P3, P4, and P5 with PBS and nitazoxanide (10 mg/kg/d or 20 mg/kg/d), using antibodies against the indicated proteins. (c, e) Densitometric analyses and quantification of (c) pS6, (e) pSMAD1/5/8 and ID1 relative protein levels. Data represent mean ± SEM (n = 4 mice per group (b) and n = 8, 7, 8 mice (d) for CTRL, NTZ 10 mg/kg/d and NTZ 20 mg/kg/d, respectively). *p<0.05, **p<0.01, ****p<0.0001, Kruskal-Wallis followed by Dunn’s multiple-comparisons test (c; e, pSMAD1/5/8) and one-way ANOVA followed by Tukey’s multiple-comparisons (e, ID1). (f) Experimental strategy of nitazoxanide treatment of BMP9/10ib mice. Pups were injected i.p. from P3 to P5 with 20 mg/kg/d nitazoxanide and on P3 and P4 with anti-BMP9 (12.5 mg/kg) and anti-BMP10 (25 mg/kg) antibodies. Arrowheads indicate the timepoints of injections and analyses. Pups were euthanized on P6 for analysis. (g-i) Representative fluorescence microscopy images of P6 retinas stained with isolectin B4, from pups injected at P3, P4, and P5 with PBS (CTRL; g), BMP9/10 blocking antibodies (h), or BMP9/10 blocking antibodies together with nitazoxanide (20 mg/kg/d; i). a, artery; v, vein. Scale bars, 500 µm. (j) AVM number and (k) AVM diameter in the retinal vasculature of pups treated as in (f). Data represent mean ± SEM (n = 5 mice per group). *p<0.05, **p<0.01, Mann-Whitney test. (l-n) Higher magnification of the region around the optic nerve showing AVMs (white arrow). Scale bars, 250 µm. (o) Retinal vein diameter of pups treated as in (f). Data represent mean ± SEM. *p<0.05, **p<0.01, one-way ANOVA followed by Tukey’s multiple-comparisons test. (p-u) Images showing retinal vasculature at the (p-r) plexus between artery and vein or (s-u) arterial vascular front of pups treated as in (f). Scale bars, (p-r) 200 µm or (s-u) 100 µm. (v, w) Quantification of vascular density at the (v) plexus and (w) arterial vascular front under control (PBS), BMP9/10 antibody (HHT), and BMP9/10 antibody plus nitazoxanide treatment. Data represent mean ± SEM. *p<0.05, **p<0.01, ***p<0.001, ****p<0.0001, (v) one-way ANOVA followed by Tukey’s multiple-comparisons test and (w) one-way ANOVA followed by Dunnett’s T3 multiple-comparisons.

### Nitazoxanide restores endothelial SMAD1/5/8 signaling in the BMP9/BMP10-immunoblocked mice

In light of the potential of nitazoxanide to simultaneously inhibit mTOR overactivation and restore SMAD1/5/8 signaling in an HHT mouse model, we examined its impact on SMAD1/5/8 signaling. For this, we took advantage of our dose-finding experiment in wild (CTRL) mice (Fig. 4d-e) and evaluated pSMAD1/5/8 and ID1 expression in the lungs of BMP9/BMP10-immunoblocked mice. The analysis of lung tissue revealed that BMP9/10 immunoblocking strongly reduced pSMAD1/5/8 and ID1 expression, both of which were restored by nitazoxanide treatment (Fig. 5a-b). Additionally, immunohistochemical (IHC) analysis confirmed increased nuclear pSMAD1/5/8 signal in CD31-positive ECs from nitazoxanide-treated lungs compared with PBS-treated controls (Fig. 5c-g). These data demonstrate that nitazoxanide restores endothelial SMAD1/5/8 signaling in the mouse lungs *in vivo*.

**Figure 5.**
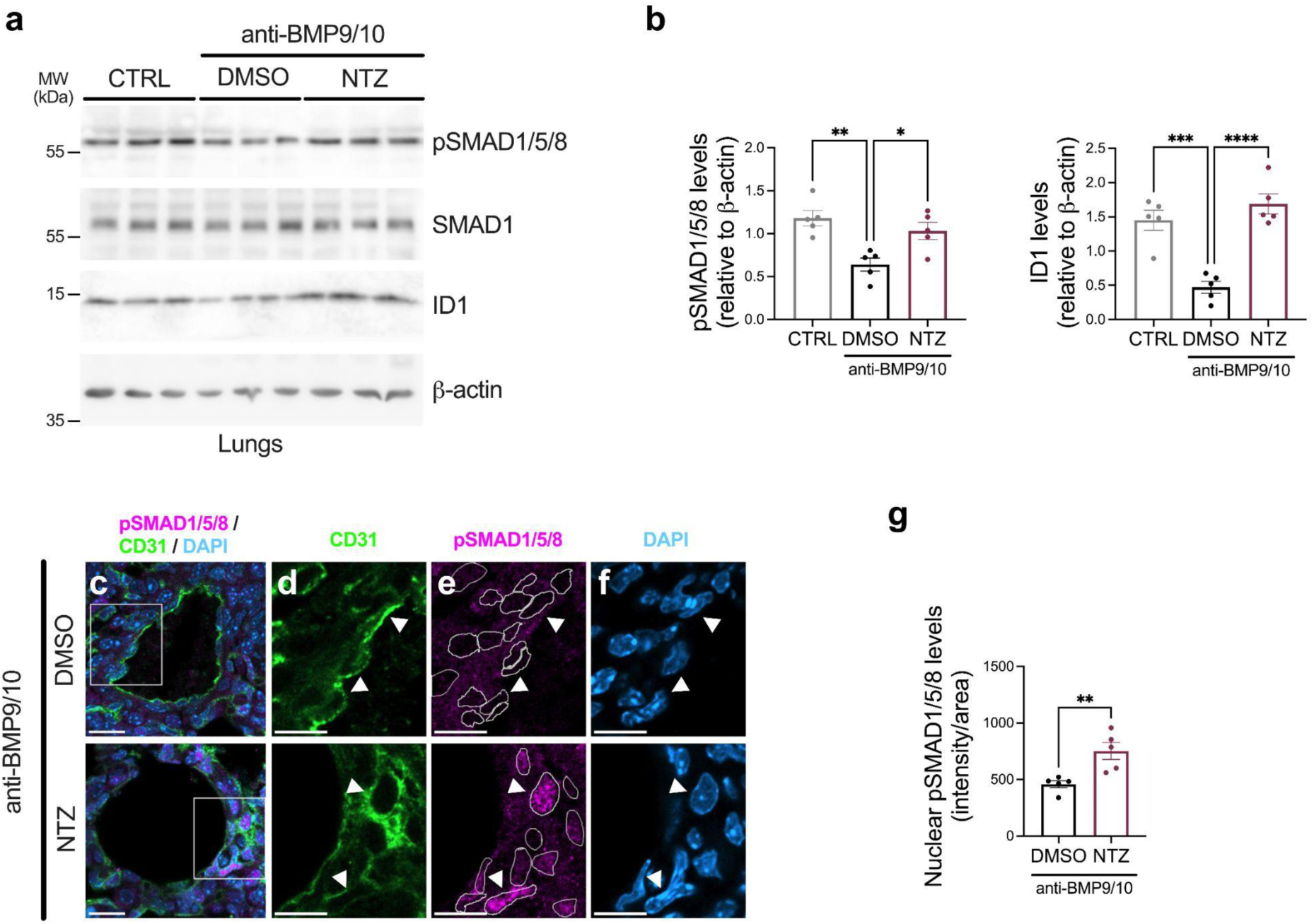
Nitazoxanide rescues SMAD1/5/8 signaling in the BMP9/10ib mice. (a) Western blot analysis of lung tissue homogenates from P6 pups injected at P3, P4, and P5 with PBS, BMP9/10 blocking antibodies, or BMP9/10 blocking antibodies plus nitazoxanide (20 mg/kg/d), using antibodies against the indicated proteins. (b) Densitometric analyses and quantification of pSMAD1/5/8 and ID1 relative levels. Data represent mean ± SEM (n = 5 mice per group). *p<0.05, **p<0.01, ***p<0.001, ****p<0.0001, one-way ANOVA followed by Tukey’s multiple-comparisons test. (c-f) Representative lungs sections from pups treated with BMP9/10 blocking antibodies (top) or BMP9/10 blocking antibodies plus nitazoxanide (bottom), stained with (d) anti-CD31 (green), (e) anti-pSMAD1/5/8 (magenta) and (f) DAPI (blue). Scale bars, 15 µm. Merged images are shown on the left (c). Scale bars, 20 µm. (g) Quantification of nuclear pSMAD1/5/8 intensity within endothelial cells. Data represent mean ± SEM (n = 5 mice per group). **p<0.01, unpaired two-tailed Student’s t-test.

### Nitazoxanide inhibits mTOR in HUVECs and prevents mTOR overactivation in vivo in BMP9/BMP10-immunoblocked mice

As previously demonstrated, mTOR signaling and phosphorylation of its downstream effector ribosomal protein S6 (S6) are increased in VEGF-stimulated ECs, and are even more pronounced in ECs within AVMs in ALK1^ΔiEC^ mouse retinas and in BMP9/10ib mice [28,32]. In all cases, these effects are completely inhibited by sirolimus, a well-known mTOR inhibitor [32].

Given that nitazoxanide has been reported to inhibit mTOR signaling in other cell types [50], we investigated whether a similar effect occurs in ECs. To this end, we compared the effects of nitazoxanide with those of sirolimus and tacrolimus (which does not inhibit mTOR) on mTOR pathway activity by assessing phosphorylation of S6. HUVECs were cultured in complete medium, conditioned for 2 days, and treated with nitazoxanide, sirolimus, or tacrolimus (1 µM) for 24 hours. Western blot analysis revealed that nitazoxanide moderately but consistently reduced S6 phosphorylation (Fig. 6a-b).

**Figure 6.**
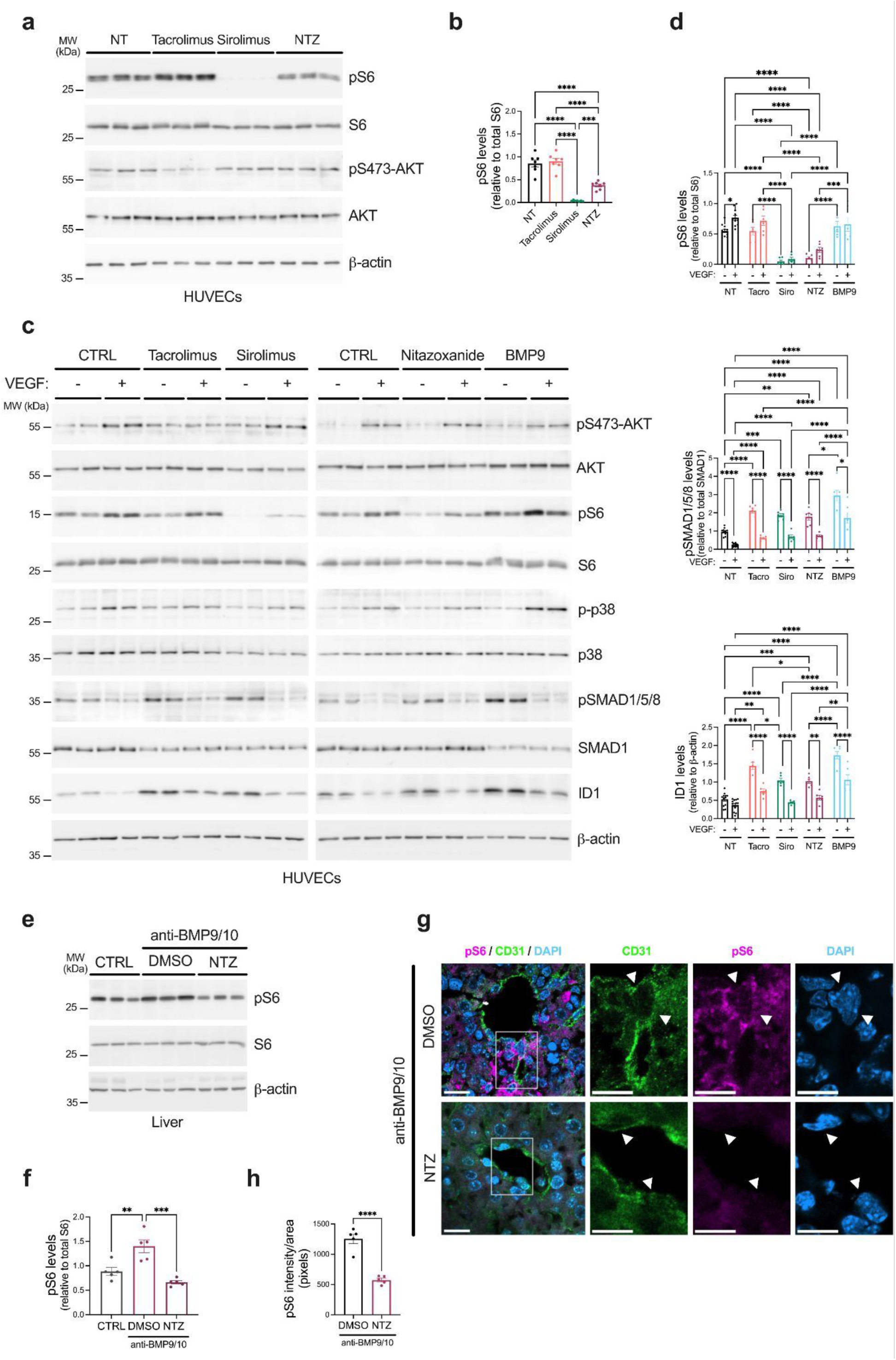
Nitazoxanide inhibits mTOR’s VEGF-mediated activation in endothelial cells and prevents mTOR overactivation in BMP9/10ib mice. (a-b) HUVECs were treated or not (NT) for 24 hours in complete medium (conditioned for 2 days) with 1 μM tacrolimus, 1μM sirolimus or 1 μM nitazoxanide. (a) Cell extracts were analyzed by Western blot using antibodies directed against the indicated proteins. (b) Densitometric analyses and quantification of pS6 relative levels in n=3 independent experiments. Data represent mean ± SEM. ***p<0.001, ****p<0.0001, one-way ANOVA followed by Tukey’s multiple-comparisons test. (c-d) HUVECs treated or not (CTRL) for 24 h with tacrolimus (1 μM), sirolimus (1 μM), nitazoxanide (1 μM) or BMP9 (10 ng/ml), were stimulated for 5 min with VEGF (25 ng/ml). (c) Cell extracts were analyzed by Western blot using antibodies directed against the indicated proteins. (d) Densitometric analyses and quantification of pS6, pSMAD1/5/8 and ID1 relative levels in n=3 independent experiments. Data represent mean ± SEM. *p<0.05, **p<0.01, ***p<0.001, ****p<0.0001, two-way ANOVA followed by Tukey’s multiple-comparisons test. For pSMAD1/5/8, data were log-transformed before analysis to meet normality and homoscedasticity test assumptions. (e) Western blot analysis of liver tissue homogenates from P6 pups injected at P3, P4, and P5 with PBS, BMP9/10 blocking antibodies, or BMP9/10 blocking antibodies plus nitazoxanide (20 mg/kg/d), using antibodies against the indicated proteins. (f) Densitometric analyses and quantification of pS6 relative levels. Data represent mean ± SEM (n = 5 mice per group). **p<0.01, ***p<0.001, one-way ANOVA followed by Tukey’s multiple-comparisons test. (g) Representative liver sections from pups treated with BMP9/10 blocking antibodies (top panels) or BMP9/10 blocking antibodies plus nitazoxanide (bottom panels), stained with anti-CD31 (green), anti-pS6 (magenta) and DAPI (blue). Scale bars, 10 µm. Merged images are shown on the left. Scale bars, 15 µm. (h) Quantification of pS6 intensity within endothelial cells. Data represent mean ± SEM (n = 5 mice per group). ****p<0.0001, unpaired two-tailed Student’s t-test.

Then, we sought to determine whether nitazoxanide is still effective in the reduction of mTOR signaling under *in vitro* conditions more relevant to angiogenesis, by performing brief VEGF stimulation in primary ECs. We found that in vehicle-pretreated (CTRL) HUVECs, VEGF stimulation robustly increased AKT and mTOR signaling, evidenced by an increase in AKT and S6 phosphorylation, whereas the BMP9-ALK1-SMAD signaling cascade remained completely blocked (Fig. 6c-d). Both pSMAD1/5/8 and ID1 levels were significantly reduced (Fig. 6c-d). On the other hand, pretreatment with BMP9 for 24 hours generated no change in AKT and mTOR signaling effect upon VEGF stimulation, while strongly activating BMP9-ALK1-SMAD signaling cascade (Fig. 6c-d). Surprisingly, nitazoxanide treatment of HUVEC for 24 hours before VEGF stimulation resulted in moderate but consistent inhibition of mTOR signaling shown by a reduction in S6 phosphorylation (Fig. 6c-d). The effect of nitazoxanide was mTOR-specific, because the drug did not affect VEGF-mediated AKT and p38 phosphorylation (Fig. 6c).

We next evaluated mTOR signaling *in vivo*. In a previous work, we reported a robust increase in mTOR signaling by assessing the level of S6 phosphorylation in whole-liver homogenates isolated from BMP9/10ib mice [32]. Our dose-finding experiments demonstrated that nitazoxanide (20 mg/kg/day) significantly reduced S6 phosphorylation in the liver of wild-type mice (Fig. 4b-c), confirming effective pathway modulation *in vivo*. Importantly, nitazoxanide treatment blocked S6 phosphorylation and normalized the signaling in the BMP9/10ib mouse livers (Fig. 6e-f). IHC analysis further revealed marked pS6 immunoreactivity in ECs of BMP9/10ib livers, which was significantly reduced following nitazoxanide treatment of the mice (Fig. 6g-h). Collectively, these data confirm that nitazoxanide could efficiently block endothelial mTOR dysregulation *in vivo*.

### Nitazoxanide activates SMAD1/5/8 signaling in endothelial cells derived from HHT patients

Finally, we evaluated the effect of nitazoxanide in a clinically relevant human model, using blood outgrowth endothelial cells (BOECs) derived from HHT patients. Cells from six patients, including individuals with ALK1 (HHT2) or SMAD4 (HHT/JPS) mutations, were treated with nitazoxanide (1 μM) for 24 hours. Under basal conditions, patient-derived BOECs exhibited reduced pSMAD1/5/8 and ID1 levels compared with healthy controls (Suppl. Fig. 4), confirming the BMP9-ALK1-SMAD signaling cascade defect. Nitazoxanide treatment significantly increased SMAD1/5/8 phosphorylation and ID1 expression in all patient samples (Fig. 7a-b), despite the presence of disease-causing mutations. These results demonstrate that nitazoxanide can activate the BMP9-ALK1-SMAD signaling in ECs derived from HHT patients, supporting its potential to restore pathway activity in a clinically relevant context.

**Figure 7.**
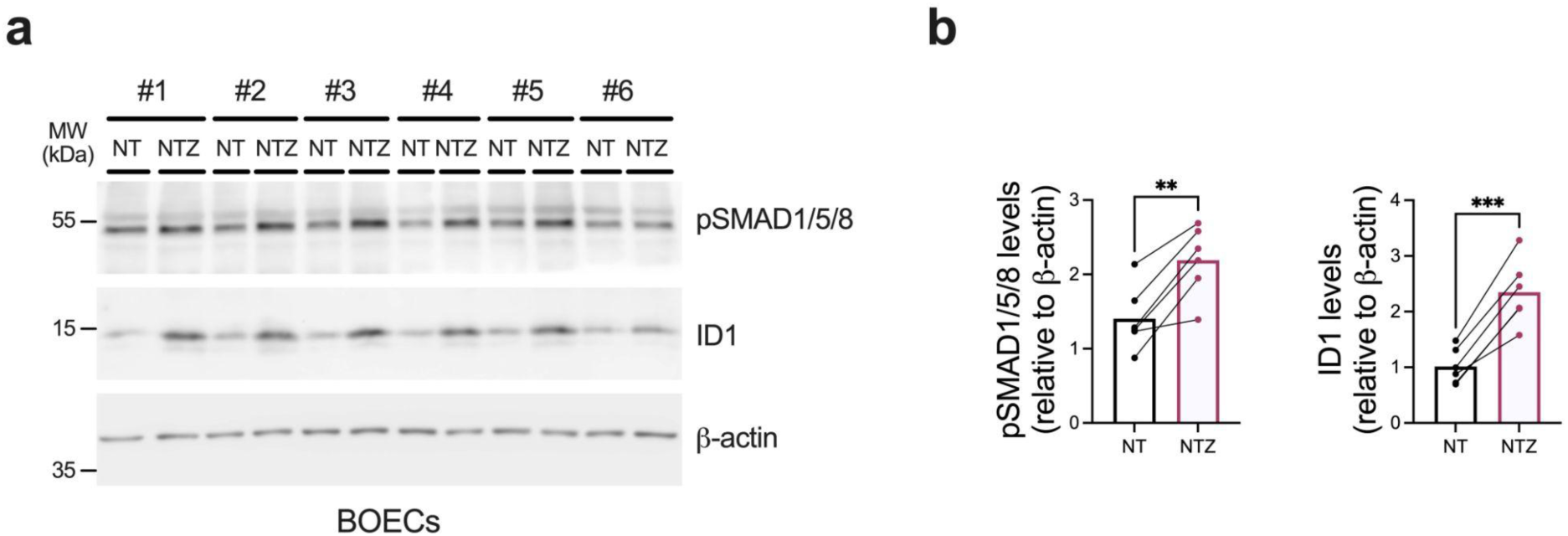
Nitazoxanide activates the BMP9-ALK1-SMAD signaling pathway in HHT patients-derived BOECs. (a) BOECs from six HHT patients–two HHT2 (ALK1-mutant; #1 and #2) and four HHT/JPS (SMAD4-mutant; #3-6)–were treated or not (NT) for 24 h with 1 μM nitazoxanide. Protein extracts were analyzed by Western blot using antibodies against the indicated proteins. (b) Densitometric analyses and quantification of pSMAD1/5/8 and ID1 relative levels. Data represent mean ± SEM. **p<0.01, ***p<0.001, paired two-tailed Student’s t-test.

## Discussion

Although HHT remains a severely debilitating disorder associated with life-threatening complications, the past decade has witnessed significant advances in therapeutic strategies targeting angiogenic cues and dysregulated intracellular pathways, substantially expanding our understanding of disease mechanisms and improving clinical management [52,53]. A growing body of preclinical research has generated diverse pharmacological and biologic candidates with clear translational potential for evaluation in clinical trials [52,53].

Among these candidates, tacrolimus has progressed along the translational pipeline, demonstrating activation of endothelial pSMAD1/5/8 signaling and amelioration of vascular pathology in experimental models of HHT [31] and PAH [53], as well as in patients [45–48]. Its structural analog, sirolimus (rapamycin), subsequently emerged as a more potent inhibitor of AVM development in HHT mouse models [32]. Notably, sirolimus was shown to exert dual modulatory activity in primary endothelial cells–including HHT patient-derived cells–and *in vivo*, simultaneously enhancing pSMAD1/5/8 signaling while suppressing mTOR activity [32]. Here, we identify nitazoxanide as a novel pharmacologic activator of endothelial SMAD1/5/8 signaling. Nitazoxanide restores the BMP9/10-ALK1-SMAD signaling cascade in cultured ECs, prevents vascular pathology in an HHT mouse model, and rescues signaling defects in HHT patient-derived BOECs. Importantly, nitazoxanide also exhibits a dual modulatory profile in primary ECs, concomitantly activating pSMAD1/5/8 signaling while inhibiting mTOR activity, thereby mirroring to some extent the mechanistic framework previously described for sirolimus.

Our data highlights that the mechanism by which nitazoxanide activates pSMAD1/5/8 signaling differs from that described for tacrolimus and sirolimus. Previous studies proposed that tacrolimus stimulates pSMAD1/5/8 signaling in ECs through the activation of the kinase domains of ALK1, ALK2 and ALK3, whereas sirolimus selectively stimulates ALK2 [31,32], in both cases by displacing the inhibitory protein FKBP12 from the kinase domain [53]. Here, we demonstrated that nitazoxanide activates endothelial pSMAD1/5/8 signaling through selective activation of ALK1. Although the precise mechanism underlying this effect–particularly whether nitazoxanide promotes dissociation of FKBP12 from the ALK1 kinase domain–remains to be elucidated, our findings would indicate that nitazoxanide could activate pSMAD1/5/8 signaling by stimulating the wild-type ALK1 allele in ECs in the HHT haploinsufficient context. This action may therefore compensate for ALK1 loss-of-function, restoring the BMP9/10-ALK1-SMAD signaling cascade.

Our investigation further revealed that nitazoxanide administration in the BMP9/10ib mice triggered a reduction in retinal hypervascularization and in the diameter of veins, and more importantly, elicited a reduction in AVMs number and diameter. Strikingly, nitazoxanide fully recovered the activation of pSMAD1/5/8 signaling in retinal EC of the BMP9/10ib model. Consistent with these findings, nitazoxanide also recovered pSMAD1/5/8 activation in the lungs and potently increased the levels of ID1 protein. Immunohistochemical analysis further indicated that this increase in pSMAD1/5/8 signaling was largely restricted to ECs within the lung vasculature, supporting an endothelial-specific restoration of this BMP signaling *in vivo*.

Current evidence indicates that the PI3K/AKT signaling pathway and several of its downstream effectors are overactivated in ECs and contribute to the development of AVMs in HHT mouse models [28,32]. Pharmacological inhibition of this signaling axis using PI3K/AKT inhibitors, as well as inhibition of the downstream effector mTOR with sirolimus, has been shown to normalize endothelial signaling *in vitro* and significantly reduce AVM formation in experimental models [28,32], supporting the contribution of PI3K-AKT-mTOR dysregulation to HHT vascular pathology.

In this context, our results show that nitazoxanide does not affect PI3K/AKT activation but moderately inhibits mTOR signaling in ECs. Importantly, although loss-of-function mutations in genes encoding components of the BMP pathway–including BMP9, ALK1, ENG, and SMAD4–are necessary to cause HHT, they are not sufficient to drive AVMs formation on their own [54]. This observation underlies the “second-hit” or “multiple-hits” triggering hypothesis, whereby additional pro-angiogenic stimuli are required for AVM development [27,55]. Among these, VEGF is recognized as a key regulator of angiogenesis and an important trigger of pathological vascular remodeling in HHT. Therefore, we investigated whether nitazoxanide modulates VEGF-dependent signaling pathways. Our data indicate that nitazoxanide exerts a clear inhibitory effect on mTOR signaling downstream of VEGF stimulation. In addition, we here show that, in the BMP9/10ib mouse liver, there is an increase in endothelial mTOR signaling evident by pS6 overactivation that is fully prevented by nitazoxanide treatments, along with an almost complete reduction of AVMs in our model. Altogether, these data suggest that nitazoxanide’s capacity to dampen this pro-angiogenic pathway may contribute to the reduction of vascular abnormalities observed in our experimental model.

Furthermore, evaluation of nitazoxanide in patient-derived blood outgrowth endothelial cells (BOECs) provided indirect evidence supporting its potential therapeutic relevance in HHT. These cells exhibited a markedly impaired BMP9-ALK1-ENG-SMAD signaling cascade, reflected by reduced levels of pSMAD1/5/8 and ID1 protein, consistent with the ALK1 or SMAD4 loss-of-function carried by these patients. This observation is in agreement with previous reports showing that, although ALK1 heterozygosity in these ECs does not significantly alter the global transcriptomic profile or acute SMAD1/5 response to BMP9 stimulation, these ALK1-mutated ECs display an approximately 50% reduction in ALK1 protein levels at the cell surface compared to control cells in basal condition (56). Strikingly, here nitazoxanide treatment restored pSMAD1/5/8 activation and increased ID1 protein levels in these BOECs, despite the underlying ALK1 or SMAD4 loss-of-function, indicating that nitazoxanide can at least partially restore the BMP9-ALK1-ENG-SMAD signaling cascade in HHT ECs.

Finally, it is important to mention that nitazoxanide is an orally bioavailable drug with extensive postmarketing experience, having been administered to more than 75 million adults and children worldwide [57]. Initially developed as an antiparasitic agent, it is currently used to treat several infectious diseases and exhibits broad-spectrum antimicrobial activity against parasites, bacteria, and certain viruses [58]. Beyond its antimicrobial effects, accumulating evidence indicates that nitazoxanide modulates multiple cellular pathways in different cell types and could exert beneficial effects across a range of pathological conditions [59,60]. In this work, our conclusion is that nitazoxanide is a potent activator of SMAD1/5/8 signaling in cells and a mouse model of HHT, including ECs derived from HHT patients. We thus propose that nitazoxanide has therapeutic potential in HHT.

## Acknowledgements

We thank OB. Nathalie Canobra for assistance with IRB paperwork and processes at the Centro Hospitalario Pereira Rossell (ASSE, Montevideo). We are grateful to the Department of Transfusional Medicine and Dr. Beatriz Boggia for their support. We acknowledge the HHT Patients Association of Uruguay and the Douglas Piquinela Foundation (Uruguay) for their support, as well as the Programa de Desarrollo de las Ciencias Básicas (PEDECIBA). We thank the core facilities of the Institut Pasteur de Montevideo, including the Advanced Bioimaging, Cell Biology, and Laboratory Animal Biotechnology Units, and in particular Gabriel Fernández and Dr. Martina Crispo for their support. This work was supported by the Agencia Nacional de Investigación e Innovación (ANII; grant FMV_1_2021_1_166595) and the Brain Vascular Malformation Consortium (BVMC; U54NS065705-YR14, BVMC Pilot Project Award). CC was supported by an ANII-SNB fellowship and the “Solidario” program (Institut Pasteur de Montevideo). SR was supported by ANII-SNI.

## Compliance with ethical standards

The authors have no relevant financial or non-financial interests to disclose. All procedures performed in this study involving human subject’s samples were in accordance with the ethical standards of the institutional and national research committees. Protocols were reviewed and approved by the Ethical Committee of the hospital Centro Hospitalario Pereira Rossell, Hospital Italiano de Buenos Aires and the Institut Pasteur de Montevideo. Additionally, Informed consents were obtained from all individual participants in the study. Clinical trial number: not applicable.

## Ethical approval

All procedures involving animals were compliant with and approved by the Institut Pasteur de Montevideo Institutional Animal Care and Use Committees (application 009-24).

---

**Supplementary Figure 1.**
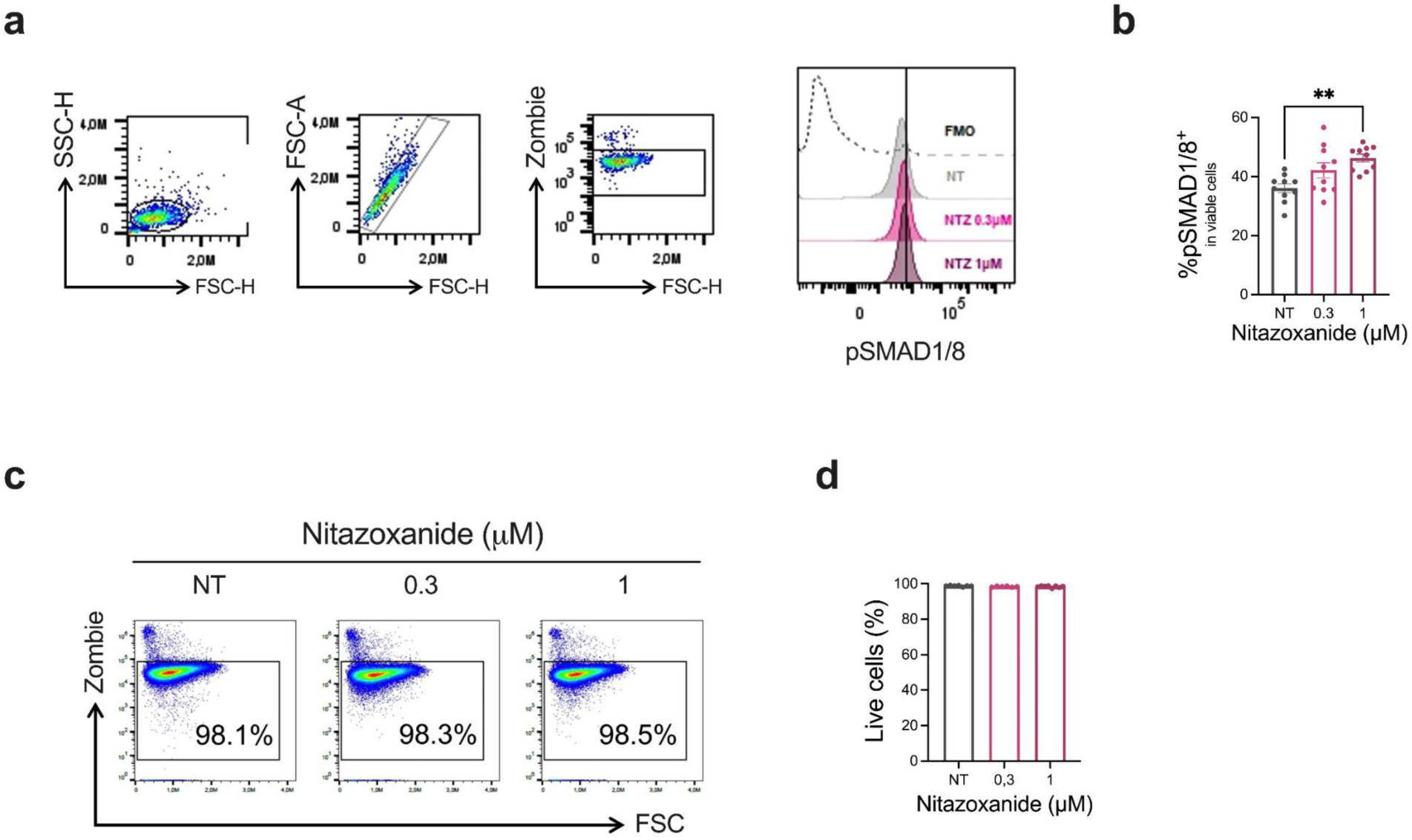
Nitazoxanide activates SMAD1/5/8 without affecting cell viability. (a-b) Flow cytometry analysis of pSMAD1/5/8 in C2C12 cells treated or not (NT) for 24 h in complete medium (conditioned for 2 days) with different concentrations of nitazoxanide (0.3 and 1 μM), in n=3 independent experiments. (a) Gating strategy used for the analysis. Cells were identified based on side scatter channel area (SSC-A) and forward scatter channel area (FSC-A), singlets were selected and dead cells were excluded using the Zombie viability marker. pSMAD1/8 positivity was determined using FMO (Fluorescence Minus One) controls in each condition and fluorescence intensity was analyzed. (b) Percentage of pSMAD1/8 positive cells. Data represent mean ± SEM. **p<0.01, one-way ANOVA followed by Tukey’s multiple-comparisons test. (c) Gating strategy for the analysis of viability of C2C12 cells treated as in (a-b). (d) Percentage of live (Zombie^−^) C2C12 cells, one-way ANOVA followed by Tukey’s multiple-comparisons test.

**Supplementary Figure 2.**
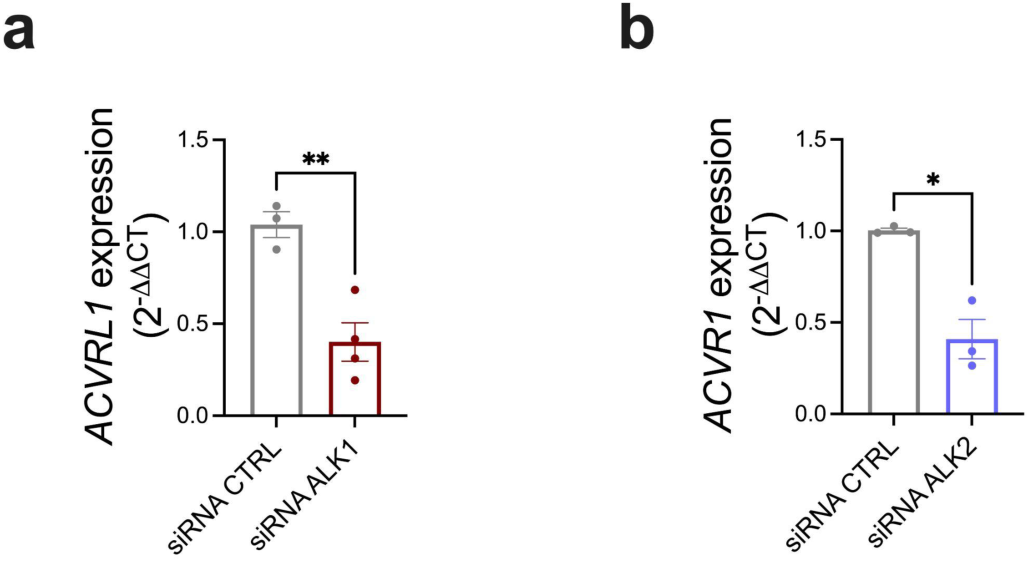
Histogram showing (a) *ACVRL1* (ALK1) and (b) *ACVR1* (ALK2) expression (2^−ΔΔCt^) in HUVECs treated with CTRL or *ACVRL1*- and *ACVR1*-targeting siRNA. Data represent mean ± SEM. *p<0.05, **p<0.01, unpaired two-tailed Student’s t-test (a) and unpaired two-tailed Student’s t-test with Welch’s correction (b).

**Supplementary Figure 3.**
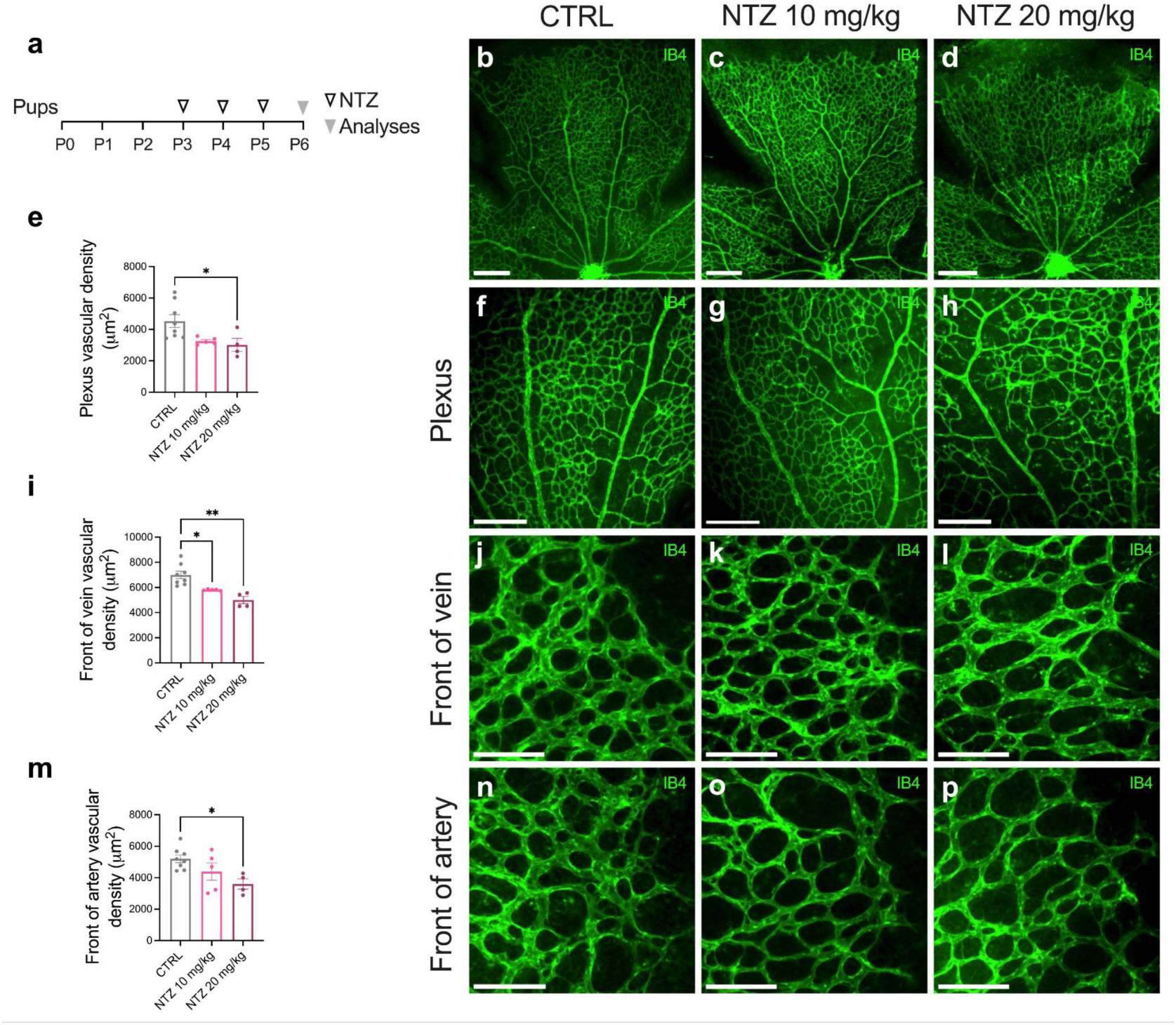
Dose-dependent effects of nitazoxanide on retinal vascular development of wild type mice. (a) Experimental strategy. Pups were injected i.p. from P3 to P5 with 10 mg/kg/d or 20 mg/kg/d nitazoxanide. Arrowheads indicate the timepoints of injections and analyses. Pups were euthanized on P6 for analysis. (b-d) Representative fluorescence microscopy images of P6 retinas stained with isolectin B4, from pups injected at P3, P4, and P5 with (b) PBS, (c) nitazoxanide 10 mg/kg/d, (d) or 20 mg/kg/d. Scale bars, 300 µm. (f-h, j-l, n-p) Representative images of the retinal vasculature showing (f-h) the plexus between artery and vein, (j-l) the venous vascular front, and (n-p) the arterial vascular front in pups treated as in (b-d). Scale bars, 200 µm (f-h) 100 µm (j-l, n-p). (e, i, m) Quantification of vascular density in the plexus (e), venous (i), and arterial (m) vascular fronts in pups treated as in (b-d). Data represent mean ± SEM (n = 4, 3, 3 mice for CTRL, NTZ 10 mg/kg/d and NTZ 20 mg/kg/d, respectively). *p<0.05, **p<0.01, one-way ANOVA followed by Tukey’s multiple-comparisons test (e, m) and one-way ANOVA followed by Dunnett’s T3 multiple-comparisons (i).

**Supplementary Figure 4.**
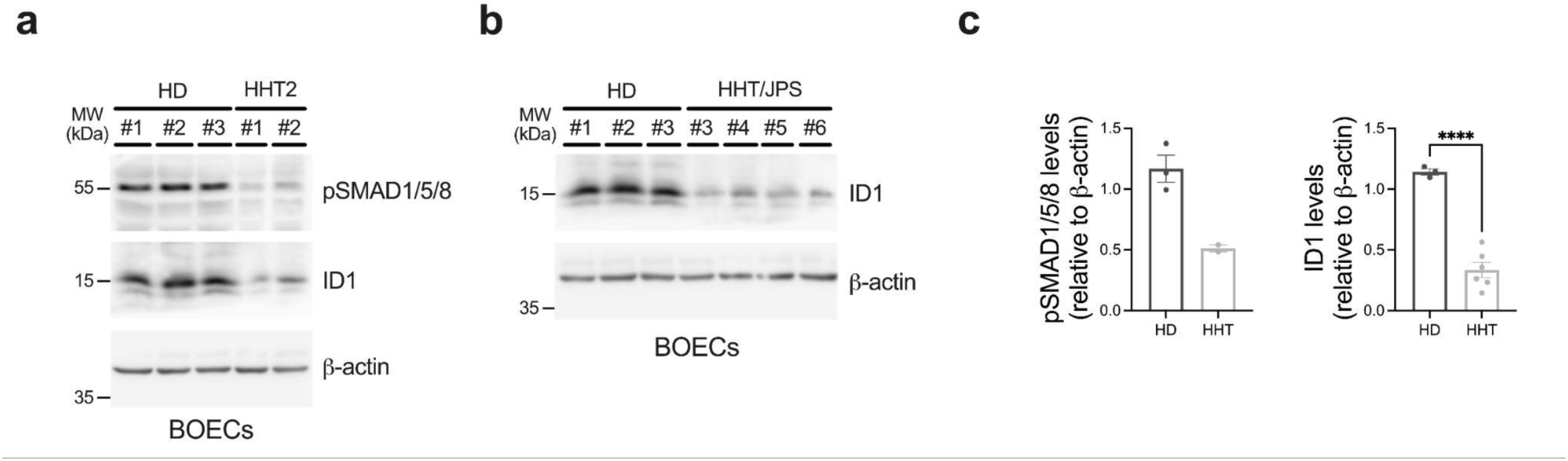
Patients derived BOECs show defects in the BMP9-ALK1-SMAD signaling cascade. (a-b) Western blot analysis of BOECs from three healthy donors and from six HHT patients: two HHT2 patients carrying mutations in ALK1 (a) and four HHT/JPS patients carrying mutations in SMAD4 (b). (c) Densitometric analyses and quantification of pSMAD1/5/8 and ID1 relative protein levels. Data represent mean ± SEM. ***p<0.001, ****p<0.0001. Statistical analyses used: unpaired two-tailed Student’s t-test.

## Notes

### Competing Interest Statement

The authors have declared no competing interest.

## References

1. Shovlin CL. Hereditary haemorrhagic telangiectasia: pathophysiology, diagnosis and treatment. Blood Rev. 2010 Nov;24(6):203–19. doi: 10.1016/j.blre.2010.07.001.

2. Faughnan ME, Mager JJ, Hetts SW, Palda VA, Lang-Robertson K, Buscarini E, Deslandres E, Kasthuri RS, Lausman A, Poetker D, et al. Second International Guidelines for the Diagnosis and Management of Hereditary Hemorrhagic Telangiectasia. Ann Intern Med. 2020 Dec 15;173(12):989–1001. doi: 10.7326/M20-1443.

3. McAllister KA, Grogg KM, Johnson DW, Gallione CJ, Baldwin MA, Jackson CE, Helmbold EA, Markel DS, McKinnon WC, Murrell J, et al. Endoglin, a TGF-beta binding protein of endothelial cells, is the gene for hereditary haemorrhagic telangiectasia type 1. Nat Genet. 1994 Dec;8(4):345–51. doi: 10.1038/ng1294-345.

4. Johnson DW, Berg JN, Baldwin MA, Gallione CJ, Marondel I, Yoon SJ, Stenzel TT, Speer M, Pericak-Vance MA, Diamond A, et al. Mutations in the activin receptor-like kinase 1 gene in hereditary haemorrhagic telangiectasia type 2. Nat Genet. 1996 Jun;13(2):189–95. doi: 10.1038/ng0696-189.

5. Wetzel-Strong SE, Detter MR, Marchuk DA. The pathobiology of vascular malformations: insights from human and model organism genetics. J Pathol. 2017 Jan;241(2):281–293. doi: 10.1002/path.4844.

6. Gallione CJ, Repetto GM, Legius E, Rustgi AK, Schelley SL, Tejpar S, Mitchell G, Drouin E, Westermann CJ, Marchuk DA. A combined syndrome of juvenile polyposis and hereditary haemorrhagic telangiectasia associated with mutations in MADH4 (SMAD4). Lancet. 2004 Mar 13;363(9412):852–9. doi: 10.1016/S0140-6736(04)15732-2.

7. Wooderchak-Donahue WL, McDonald J, O’Fallon B, Upton PD, Li W, Roman BL, Young S, Plant P, Fülöp GT, Langa C, et al. BMP9 mutations cause a vascular-anomaly syndrome with phenotypic overlap with hereditary hemorrhagic telangiectasia. Am J Hum Genet. 2013 Sep 5;93(3):530–7. doi: 10.1016/j.ajhg.2013.07.004.

8. Goumans MJ, Zwijsen A, Ten Dijke P, Bailly S. Bone Morphogenetic Proteins in Vascular Homeostasis and Disease. Cold Spring Harb Perspect Biol. 2018 Feb 1;10(2):a031989. doi: 10.1101/cshperspect.a031989.

9. Akhurst RJ, Hata A. Targeting the TGFβ signalling pathway in disease. Nat Rev Drug Discov. 2012 Oct;11(10):790–811. doi: 10.1038/nrd3810.

10. Newman JH, Trembath RC, Morse JA, Grunig E, Loyd JE, Adnot S, Coccolo F, Ventura C, Phillips JA 3rd, Knowles JA, et al. Genetic basis of pulmonary arterial hypertension: current understanding and future directions. J Am Coll Cardiol. 2004 Jun 16;43(12 Suppl S):33S–39S. doi: 10.1016/j.jacc.2004.02.028.

11. Scharpfenecker M, van Dinther M, Liu Z, van Bezooijen RL, Zhao Q, Pukac L, Löwik CW, ten Dijke P. BMP-9 signals via ALK1 and inhibits bFGF-induced endothelial cell proliferation and VEGF-stimulated angiogenesis. J Cell Sci. 2007 Mar 15;120(Pt 6):964–72. doi: 10.1242/jcs.002949.

12. David L, Mallet C, Mazerbourg S, Feige JJ, Bailly S. Identification of BMP9 and BMP10 as functional activators of the orphan activin receptor-like kinase 1 (ALK1) in endothelial cells. Blood. 2007 Mar 1;109(5):1953–61. doi: 10.1182/blood-2006-07-034124.

13. Brown MA, Zhao Q, Baker KA, Naik C, Chen C, Pukac L, Singh M, Tsareva T, Parice Y, Mahoney A, et al. Crystal structure of BMP-9 and functional interactions with pro-region and receptors. J Biol Chem. 2005 Jul 1;280(26):25111–8. doi: 10.1074/jbc.M503328200.

14. Salmon RM, Guo J, Wood JH, Tong Z, Beech JS, Lawera A, Yu M, Grainger DJ, Reckless J, Morrell NW, et al. Molecular basis of ALK1-mediated signalling by BMP9/BMP10 and their prodomain-bound forms. Nat Commun. 2020 Apr 1;11(1):1621. doi: 10.1038/s41467-020-15425-3.

15. Ricard N, Bailly S, Guignabert C, Simons M. The quiescent endothelium: signalling pathways regulating organ-specific endothelial normalcy. Nat Rev Cardiol. 2021 Aug;18(8):565–580. doi: 10.1038/s41569-021-00517-4.

16. Massagué J. TGFβ signalling in context. Nat Rev Mol Cell Biol. 2012 Oct;13(10):616–30. doi: 10.1038/nrm3434.

17. Cai J, Pardali E, Sánchez-Duffhues G, ten Dijke P. BMP signaling in vascular diseases. FEBS Lett. 2012 Jul 4;586(14):1993–2002. doi: 10.1016/j.febslet.2012.04.030.

18. Alaa El Din F, Patri S, Thoreau V, Rodriguez-Ballesteros M, Hamade E, Bailly S, Gilbert-Dussardier B, Abou Merhi R, et al. Functional and splicing defect analysis of 23 ACVRL1 mutations in a cohort of patients affected by Hereditary Hemorrhagic Telangiectasia. PLoS One. 2015 Jul 15;10(7):e0132111. doi: 10.1371/journal.pone.0132111.

19. Ricard N, Bidart M, Mallet C, Lesca G, Giraud S, Prudent R, Feige JJ, Bailly S. Functional analysis of the BMP9 response of ALK1 mutants from HHT2 patients: a diagnostic tool for novel ACVRL1 mutations. Blood. 2010 Sep 2;116(9):1604–12. doi: 10.1182/blood-2010-03-276881.

20. Ruiz-Llorente L, Gallardo-Vara E, Rossi E, Smadja DM, Botella LM, Bernabeu C. Endoglin and alk1 as therapeutic targets for hereditary hemorrhagic telangiectasia. Expert Opin Ther Targets. 2017 Oct;21(10):933–947. doi: 10.1080/14728222.2017.1365839.

21. Tual-Chalot S, Oh SP, Arthur HM. Mouse models of hereditary hemorrhagic telangiectasia: recent advances and future challenges. Front Genet. 2015 Feb 18;6:25. doi: 10.3389/fgene.2015.00025.

22. Ricard N, Ciais D, Levet S, Subileau M, Mallet C, Zimmers TA, Lee SJ, Bidart M, Feige JJ, Bailly S. BMP9 and BMP10 are critical for postnatal retinal vascular remodeling. Blood. 2012 Jun 21;119(25):6162–71. doi: 10.1182/blood-2012-01-407593.

23. Chen H, Brady Ridgway J, Sai T, Lai J, Warming S, Chen H, Roose-Girma M, Zhang G, Shou W, Yan M. Context-dependent signaling defines roles of BMP9 and BMP10 in embryonic and postnatal development. Proc Natl Acad Sci U S A. 2013 Jul 16;110(29):11887–92. doi: 10.1073/pnas.1306074110.

24. Larrivée B, Prahst C, Gordon E, del Toro R, Mathivet T, Duarte A, Simons M, Eichmann A. ALK1 signaling inhibits angiogenesis by cooperating with the Notch pathway. Dev Cell. 2012 Mar 13;22(3):489–500. doi: 10.1016/j.devcel.2012.02.005.

25. Ardelean DS, Letarte M. Anti-angiogenic therapeutic strategies in hereditary hemorrhagic telangiectasia. Front Genet. 2015 Feb 11;6:35. doi: 10.3389/fgene.2015.00035.

26. Thalgott J, Dos-Santos-Luis D, Lebrin F. Pericytes as targets in hereditary hemorrhagic telangiectasia. Front Genet. 2015 Feb 13;6:37. doi: 10.3389/fgene.2015.00037.

27. Arthur HM, Roman BL. An update on preclinical models of hereditary haemorrhagic telangiectasia: Insights into disease mechanisms. Front Med (Lausanne). 2022 Sep 29;9:973964. doi: 10.3389/fmed.2022.973964.

28. Ola R, Dubrac A, Han J, Zhang F, Fang JS, Larrivée B, Lee M, Urarte AA, Kraehling JR, Genet G, et al. PI3 kinase inhibition improves vascular malformations in mouse models of hereditary haemorrhagic telangiectasia. Nat Commun. 2016 Nov 29;7:13650. doi: 10.1038/ncomms13650.

29. Jin Y, Muhl L, Burmakin M, Wang Y, Duchez AC, Betsholtz C, Arthur HM, Jakobsson L. Endoglin prevents vascular malformation by regulating flow-induced cell migration and specification through VEGFR2 signalling. Nat Cell Biol. 2017 Jun;19(6):639–652. doi: 10.1038/ncb3534.

30. Kim JH, Peacock MR, George SC, Hughes CC. BMP9 induces EphrinB2 expression in endothelial cells through an Alk1-BMPRII/ActRII-ID1/ID3-dependent pathway: implications for hereditary hemorrhagic telangiectasia type II. Angiogenesis. 2012 Sep;15(3):497–509. doi: 10.1007/s10456-012-9277-x.

31. Ruiz S, Chandakkar P, Zhao H, Papoin J, Chatterjee PK, Christen E, Metz CN, Blanc L, Campagne F, Marambaud P. Tacrolimus rescues the signaling and gene expression signature of endothelial ALK1 loss-of-function and improves HHT vascular pathology. Hum Mol Genet. 2017 Dec 15;26(24):4786–4798. doi: 10.1093/hmg/ddx358.

32. Ruiz S, Zhao H, Chandakkar P, Papoin J, Choi H, Nomura-Kitabayashi A, Patel R, Gillen M, Diao L, Chatterjee PK, et al. Correcting Smad1/5/8, mTOR, and VEGFR2 treats pathology in hereditary hemorrhagic telangiectasia models. J Clin Invest. 2020 Feb 3;130(2):942–957. doi: 10.1172/JCI127425.

33. Walker EJ, Su H, Shen F, Degos V, Amend G, Jun K, Young WL. Bevacizumab attenuates VEGF-induced angiogenesis and vascular malformations in the adult mouse brain. Stroke. 2012 Jul;43(7):1925–30. doi: 10.1161/STROKEAHA.111.647982.

34. Banerjee K, Lin Y, Gahn J, Cordero J, Gupta P, Mohamed I, Graupera M, Dobreva G, Schwartz MA, Ola R. SMAD4 maintains the fluid shear stress set point to protect against arterial-venous malformations. J Clin Invest. 2023 Sep 15;133(18):e168352. doi: 10.1172/JCI168352.

35. Kim YH, Kim MJ, Choe SW, Sprecher D, Lee YJ, P Oh S. Selective effects of oral antiangiogenic tyrosine kinase inhibitors on an animal model of hereditary hemorrhagic telangiectasia. J Thromb Haemost. 2017 Jun;15(6):1095–1102. doi: 10.1111/jth.13683.

36. Dinakaran S, Qutaina S, Zhao H, Tang Y, Wang Z, Ruiz S, Nomura-Kitabayashi A, Metz CN, Arthur HM, Meadows SM, et al. CDK6-mediated endothelial cell cycle acceleration drives arteriovenous malformations in hereditary hemorrhagic telangiectasia. Nat Cardiovasc Res. 2024 Nov;3(11):1301–1317. doi: 10.1038/s44161-024-00550-9.

37. Gaillard S, Dupuis-Girod S, Boutitie F, Rivière S, Morinière S, Hatron PY, Manfredi G, Kaminsky P, Capitaine AL, Roy P, et al. Tranexamic acid for epistaxis in hereditary hemorrhagic telangiectasia patients: a European cross-over controlled trial in a rare disease. J Thromb Haemost. 2014 Sep;12(9):1494–502. doi: 10.1111/jth.12654.

38. Faughnan ME, Gossage JR, Chakinala MM, Oh SP, Kasthuri R, Hughes CCW, McWilliams JP, Parambil JG, Vozoris N, Donaldson J, et al. Pazopanib may reduce bleeding in hereditary hemorrhagic telangiectasia. Angiogenesis. 2019 Feb;22(1):145–155. doi: 10.1007/s10456-018-9646-1.

39. Dupuis-Girod S, Rivière S, Lavigne C, Fargeton AE, Gilbert-Dussardier B, Grobost V, Leguy-Seguin V, Maillard H, Mohamed S, Decullier E, et al. Efficacy and safety of intravenous bevacizumab on severe bleeding associated with hemorrhagic hereditary telangiectasia: A national, randomized multicenter trial. J Intern Med. 2023 Dec;294(6):761–774. doi: 10.1111/joim.13714.

40. Al-Samkari H, Kasthuri RS, Iyer VN, Pishko AM, Decker JE, Weiss CR, Whitehead KJ, Conrad MB, Zumberg MS, Zhou JY, et al. Pomalidomide for Epistaxis in Hereditary Hemorrhagic Telangiectasia. N Engl J Med. 2024 Sep 19;391(11):1015–1027. doi: 10.1056/NEJMoa2312749.

41. Al-Samkari H, Hessels J, Riera-Mestre A, Dupuis-Girod S, Van Zele T, Gómez Del Olmo V, Hodges PG, Torres-Iglesias R, Bertè R, et al. Engasertib versus Placebo for Bleeding in Hereditary Hemorrhagic Telangiectasia. N Engl J Med. 2025 Nov 27;393(21):2131–2141. doi: 10.1056/NEJMoa2504411.

42. Zhang E, Kasthuri RS, Parambil J, Prasad V, Iyer VN, Whitehead KJ, Hodges PG, Pishko AM, Conrad MB, Phelan D, et al. Pomalidomide for hereditary hemorrhagic telangiectasia: after trial longitudinal assessment study (PATH-HHT ATLAS). Blood Adv. 2026 Mar 10;10(5):1799–1808. doi: 10.1182/bloodadvances.2025018382.

43. Ormiston ML, Toshner MR, Kiskin FN, Huang CJ, Groves E, Morrell NW, Rana AA. Generation and Culture of Blood Outgrowth Endothelial Cells from Human Peripheral Blood. J Vis Exp. 2015 Dec 23;(106):e53384. doi: 10.3791/53384.

44. Żyżyńska-Granica B, Koziak K. Identification of suitable reference genes for real-time PCR analysis of statin-treated human umbilical vein endothelial cells. PLoS One. 2012;7(12):e51547. doi: 10.1371/journal.pone.0051547.

45. Sommer N, Droege F, Gamen KE, Geisthoff U, Gall H, Tello K, Richter MJ, Deubner LM, Schmiedel R, Hecker M, et al. Treatment with low-dose tacrolimus inhibits bleeding complications in a patient with hereditary hemorrhagic telangiectasia and pulmonary arterial hypertension. Pulm Circ. 2019 Apr-Jun;9(2):2045894018805406. doi: 10.1177/2045894018805406.

46. Dupuis-Girod S, Fargeton AE, Grobost V, Rivière S, Beaudoin M, Decullier E, Bernard L, Bréant V, Colombet B, Philouze P, et al. Efficacy and Safety of a 0.1% Tacrolimus Nasal Ointment as a Treatment for Epistaxis in Hereditary Hemorrhagic Telangiectasia: A Double-Blind, Randomized, Placebo-Controlled, Multicenter Trial. J Clin Med. 2020 Apr 26;9(5):1262. doi: 10.3390/jcm9051262.

47. Hessels J, Kroon S, Boerman S, Nelissen RC, Grutters JC, Snijder RJ, Lebrin F, Post MC, Mummery CL, Mager JJ. Efficacy and Safety of Tacrolimus as Treatment for Bleeding Caused by Hereditary Hemorrhagic Telangiectasia: An Open-Label, Pilot Study. J Clin Med. 2022 Sep 7;11(18):5280. doi: 10.3390/jcm11185280.

48. Álvarez-Hernández P, Patier JL, Marcos S, Gómez Del Olmo V, Lorente-Herraiz L, Recio-Poveda L, Botella LM, Viteri-Noël A, Albiñana V. Tacrolimus as a Promising Drug for Epistaxis and Gastrointestinal Bleeding in HHT. J Clin Med. 2023 Nov 29;12(23):7410. doi: 10.3390/jcm12237410.

49. Droege F, Guilhem A, Ricard N, Spiekerkoetter E, Hermann R, Rossi E, Bailly S, Dupuis-Girod S, Clancy M, Friday C. Executive summary of the 15th HHT international scientific conference. Angiogenesis. 2025 Nov 18;28(Suppl 1):61. doi: 10.1007/s10456-025-09997-1.

50. Ruiz S, Zhao H, Chandakkar P, Chatterjee PK, Papoin J, Blanc L, Metz CN, Campagne F, Marambaud P. A mouse model of hereditary hemorrhagic telangiectasia generated by transmammary-delivered immunoblocking of BMP9 and BMP10. Sci Rep. 2016 Nov 22;5:37366. doi: 10.1038/srep37366.

51. Lam KK, Zheng X, Forestieri R, Balgi AD, Nodwell M, Vollett S, Anderson HJ, Andersen RJ, Av-Gay Y, Roberge M. Nitazoxanide stimulates autophagy and inhibits mTORC1 signaling and intracellular proliferation of Mycobacterium tuberculosis. PLoS Pathog. 2012;8(5):e1002691. doi: 10.1371/journal.ppat.1002691.

52. Al Tabosh T, Al Tarrass M, Tourvieilhe L, Guilhem A, Dupuis-Girod S, Bailly S. Hereditary hemorrhagic telangiectasia: from signaling insights to therapeutic advances. J Clin Invest. 2024 Feb 15;134(4):e176379. doi: 10.1172/JCI176379.

53. Ficany A, Del Alamo M, Bernabeu C, Shovlin CL, Rossi E. Epistaxis Prevention, Treatment, and Future Perspectives for Hereditary Hemorrhagic Telangiectasia. J Clin Med. 2025 Oct 30;14(21):7724. doi: 10.3390/jcm14217724.

54. Bernabeu C, Bayrak-Toydemir P, McDonald J, Letarte M. Potential Second-Hits in Hereditary Hemorrhagic Telangiectasia. J Clin Med. 2020 Nov 5;9(11):3571. doi: 10.3390/jcm9113571.

55. Spiekerkoetter E, Tian X, Cai J, Hopper RK, Sudheendra D, Li CG, El-Bizri N, Sawada H, Haghighat R, Chan R, et al. FK506 activates BMPR2, rescues endothelial dysfunction, and reverses pulmonary hypertension. J Clin Invest. 2013 Aug;123(8):3600–13. doi: 10.1172/JCI65592.

56. Al Tabosh T, Liu H, Koça D, Al Tarrass M, Tu L, Giraud S, Delagrange L, Beaudoin M, Rivière S, Grobost V, et al. Impact of heterozygous ALK1 mutations on the transcriptomic response to BMP9 and BMP10 in endothelial cells from hereditary hemorrhagic telangiectasia and pulmonary arterial hypertension donors. Angiogenesis. 2024 May;27(2):211–227. doi: 10.1007/s10456-023-09902-8.

57. Rossignol JF. Nitazoxanide: a first-in-class broad-spectrum antiviral agent. Antiviral Res. 2014 Oct;110:94–103. doi: 10.1016/j.antiviral.2014.07.014.

58. Walker LE, FitzGerald R, Saunders G, Lyon R, Fisher M, Martin K, Eberhart I, Woods C, Ewings S, Hale C, et al. An Open Label, Adaptive, Phase 1 Trial of High-Dose Oral Nitazoxanide in Healthy Volunteers: An Antiviral Candidate for SARS-CoV-2. Clin Pharmacol Ther. 2022 Mar;111(3):585–594. doi: 10.1002/cpt.2463.

59. Chen XY, Dong YC, Yu YY, Jiang M, Bu WJ, Li P, Sun ZJ, Dong DL. Anthelmintic nitazoxanide protects against experimental pulmonary fibrosis. Br J Pharmacol. 2023 Dec;180(23):3008–3023. doi: 10.1111/bph.16190.

60. Zhu HT, Luo J, Peng Y, Cheng XF, Wu SZ, Zhao YD, Chang L, Sun ZJ, Dong DL. Nitazoxanide protects against experimental ulcerative colitis through improving intestinal barrier and inhibiting inflammation. Chem Biol Interact. 2024 May 25;395:111013. doi: 10.1016/j.cbi.2024.111013.

